# First responders shape a prompt and sharp NF-κB–mediated transcriptional response to TNF-α

**DOI:** 10.1101/2020.06.27.174995

**Authors:** Samuel Zambrano, Alessia Loffreda, Elena Carelli, Giacomo Stefanelli, Federica Colombo, Edouard Bertrand, Carlo Tacchetti, Alessandra Agresti, Marco E. Bianchi, Nacho Molina, Davide Mazza

## Abstract

NF-κB acts as the master regulator of the transcriptional response to inflammatory signals by translocating into the nucleus upon stimuli, but we lack a single-cell characterization of the resulting transcription dynamics. Here we show that transcription of NF-κB target genes is strongly heterogeneous in individual cells but dynamically coordinated at the population level, since the average nascent transcription is prompt (i.e. occurs almost immediately) and sharp (i.e. increases and decreases rapidly) compared to NF-κB nuclear localization. Using an NF-κB-controlled MS2 reporter we confirm that the population-level transcriptional activity emerges from a strongly heterogeneous response in single cells as compared to NF-κB translocation dynamics, including the presence of a fraction of “first responders”. Mathematical models show that a combination of NF-κB mediated gene activation and a gene activity module including a gene refractory state is enough to produce sharp and prompt transcriptional responses. Our data and models show how the expression of the target genes of a paradigmatic inducible transcription activator upon stimuli can be time-resolved at population level and yet heterogeneous across single cells.

## Introduction

A tight control of gene expression is assumed to be fundamental for any living system, from prokaryotes to higher organisms. For this reason, it was surprising to find that the same gene within a clonal population of identical cells can be translated into different protein levels (Ko et al., 1990) which can fluctuate in time even within the same cell (Elowitz et al., 2002). The development of accurate techniques allowing to measure gene expression in single living cells showed that such variability is related to discontinuous transcriptional “bursts” (Tunnacliffe and Chubb, 2020), spurts of RNA production interspersed with periods of no activity, that emerge from fluctuations of the gene between “active” and “inactive” states, whose precise origin is only partially understood (Chong et al., 2014).

Transcriptional bursts have been observed for a variety of organisms(Golding et al., 2005; Pichon et al., 2018; Suter et al., 2011), but their functional role is also unclear, although it has been proposed as a natural mechanism exploited and controlled by cells to either produce variability or robustness in gene-expression programs, presumably in a context-specific way (Raj and van Oudenaarden, 2008). Transcriptional bursts are indeed modulated by external stimuli (Molina et al., 2013), by the developmental stage of the organism (Muramoto et al., 2012) and by chromatin state(Nicolas et al., 2018). However, we are still far from having a complete picture of how the delicate balance between robust control and variability in gene expression is achieved (Raj and van Oudenaarden, 2008).

Such balance is presumably gene and cell specific, and different for different biological processes. For example, the inflammatory response is characterized by a variable degree of transcriptional heterogeneity across genes, species and cell types (Hagai et al., 2018), whose connection to the dynamics of transcriptional bursting is unexplored. Transcription in inflammation depends on the dynamics of its master regulator (Hayden and Ghosh, 2008): the transcription factor NF-κB. NF-κB dimers containing the monomer p65 (that we refer to as NF-κB in what follows) are activated by re-localizing from the cytoplasm to the nucleus upon inflammatory stimuli such as tumor necrosis factor alpha (TNF-α). This activation by nuclear localization is tightly regulated by a system of negative feedbacks (Hoffmann et al., 2002) so that cells display a variety of nuclear localization dynamics of NF-κB, including oscillations (Nelson et al., 2004; Tay et al., 2010; Zambrano et al., 2014a). Population-level measurements have shown that NF-κB dynamics lead to different dynamical patterns of mRNA expression (Ashall et al., 2009; Nelson et al., 2004; Sung et al., 2009; Zambrano et al., 2016). The NF-κB mediated nascent transcriptional response to stimuli at the population level is however fast, comparable with the translocation dynamics of NF-κB (Hao and Baltimore, 2013; Zambrano et al., 2016) that peaks at 30 min–1 h depending on the cell line and is accompanied by a fast binding of NF-κB to the promoter of target genes (Saccani et al., 2001).

Much less is known about how NF-κB dynamics modulates transcriptional variability at single cell level. Time-lapse analysis of NF-κB translocation, followed by analysis of mRNA expression at a single time-point through RNA FISH (Lee et al., 2014) and scRNA-seq (Lane et al., 2017) has demonstrated that different NF-κB dynamics translate into specific gene expression programs in single cells. Direct simultaneous observation of NF-κB dynamics and its gene expression products has so far been carried out at the protein level only, using GFP-transgenes (Nelson et al., 2004). More recent studies have begun to interrogate systematically how the NF-κB mediated transcriptional dynamics is modulated at the single-cell level by making use of a destabilized GFP transgene under the control of an HIV-LTR promoter (carrying two binding sites for NF-κB (Stroud et al., 2009)). In these studies, TNF-α induced gene expression has been shown to occur in bursts that are tuned by the insertion site of the transgene (Dar et al., 2012) and that are amplified by TAT-mediated positive feedbacks upon viral activation (Wong et al., 2018). However, as these assays are based on protein reporters with limited temporal resolution, the relationship between NF-κB nuclear localization and transcriptional dynamics at single cell level and its connection with the population level remains unexplored.

To address this, here we analyzed the cellular response to TNF-α at single-cell level in terms of NF-κB localization and nascent transcription, both for multiple genes in fixed cells (by single-molecule RNA FISH) and for a MS2 reporter gene controlled by an HIV-LTR promoter (Tantale et al., 2016) in living cells (by time-lapse imaging). We find that although different genes are expressed with different degrees of variability, they share common average population dynamics of nascent transcription that is *prompt* (i.e. occurs simultaneously with NF-κB translocation) and *sharp* (i.e. it is limited in time and decays faster than NF-κB nuclear localization). Live-cell analysis combined with repeated stimulation in microfluidics reveals that the population’s sharp response is due to two factors: (i) a fraction of cells – first responders – that respond promptly and synchronously to TNF-α and (ii) a characteristic gene inactive time, during which the gene is insensitive to reactivation, following each active period. Mathematical modelling shows that indeed only the combination of transcriptional activity driven by NF-κB localization and a gene activity module including a refractory state can recapitulate the promptness and the sharpness of the transcriptional response.

Our results show how the interaction of NF-κB localization dynamics and target gene activity can produce a timely and gene-specific collective response upon inflammatory stimuli.

## Results

### Population-level NF-κB-mediated transcription is prompt and sharp, despite being heterogeneous in single cells

To characterize transcriptional dynamics of inflammatory genes at single-cell level, HeLa cells were exposed to TNF-α and mature and nascent transcripts of three (Rabani et al., 2011; Sung et al., 2009; Zambrano et al., 2016) NF-κB target genes (*NFKBIA* coding for NF-κB main inhibitor IκBα, *IL6* for the cytokine IL6 and *TNF* for the cytokine TNF-α) at different timepoints (**Figure S1A**) were quantified using single molecule fluorescence in-situ hybridization (Tsanov et al., 2016) (smFISH, see **Methods**). smFISH allows counting both nascent RNA molecules at active transcription sites (TS), which appear as 1 or 2 bright dots in the nucleus (**Figure 1A** and **Figure S1B**), and mature mRNA molecules, which appear as individual dots scattered in the nucleus and in the cytoplasm (**Figure 1A** and **Figure S1B**). In response to 10 ng/ml TNF-α, transcription of the three tested genes was induced with different degrees of cell-to-cell variability (**Figure 1A and 1B**). Such variability is captured by the Gini coefficient(Shaffer et al., 2017), a metric that ranges between 0 -when all cells express the same number of mRNAs- and 1 -when all mRNAs are detected in just one cell. *NFKBIA* displayed the most uniform expression (Gini ranging between 0.21 and 0.26, comparable to what previously reported for housekeeping genes(Shaffer et al., 2017)), while *IL6* (Gini from 0.41 to 0.55) and *TNF* (Gini from 0.29 to 0.33) were more unevenly expressed. Such different degrees of heterogeneity of the analyzed genes can be related to different bursting kinetics (Tunnacliffe and Chubb, 2020). By fitting the distribution of mature RNAs in single cells to a simple negative binomial model (Tunnacliffe and Chubb, 2020) whose parameters depend on the bursts’ features (Raj et al., 2006) (**Figure S1C**) we estimate a higher relative burst frequency for *TNF* and *NFKBIA* than for *IL6*. The gene activity at single-cell level, estimated as the fractions of cells carrying active TS, indeed strongly differed among the genes considered: after stimulation *NFKBIA* TS were detectable in the largest fraction of cells (ranging from 84% at 20 min to 44% at 3h post TNF-α) followed by *IL6*TS (ranging from 32% to 21%) and *TNF* TS (from 16% to 9%).

**Figure 1.**
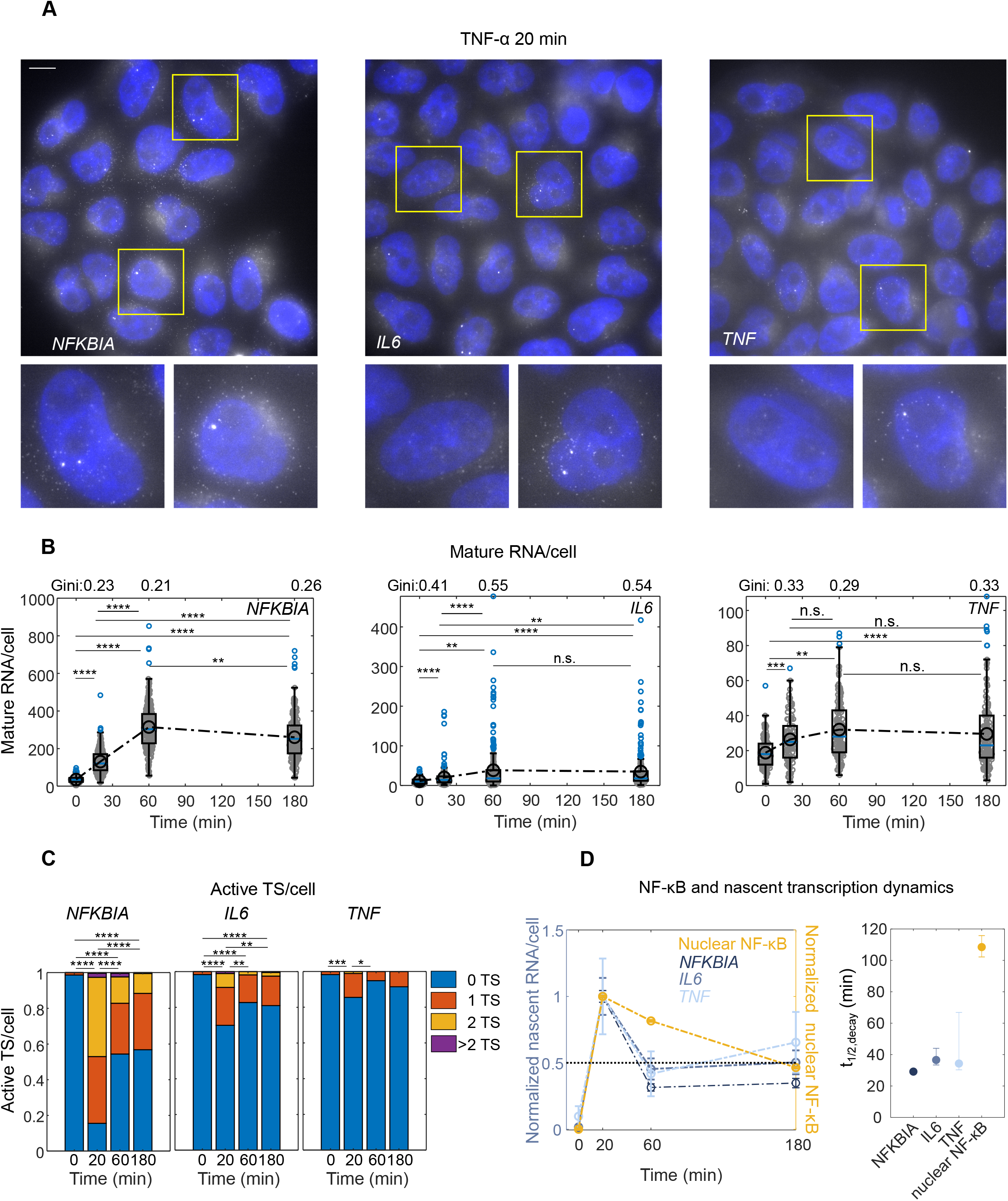
Nascent transcription of NF-κB target genes is prompt and sharp. **A.** Exemplary smFISH acquisitions using probes targeting *NFKBIA, IL6* and *TNF* RNAs 20 min after induction with TNF-α. Maximum projection, scale bar 10 μm. **B.** Mature *MS2* transcripts per cell measured at different times following TNF-α. Also displayed is the Gini coefficient measured at 20’, 1 hour and 3 hour after stimulation, as an estimate of the heterogeneity in the cell-by-cell expression of the three targets (*n_cells_* =219, 270, 250, 206 for 0’, 20’, 1 hour, 3 hours; *IL6*: *n_cells_* =187, 193, 220, 220 for 0’, 20’, 1 hour, 3 hours; *TNF*: *n_cells_* =117, 90, 157, 140 Kruskal-Wallis test (KW), * p<0.05, ** p< 0.01, *** p<0.001, **** p<0.0001). **C.** Fraction of cells with either 0,1,2 or >2 active transcription sites for MS2 measured by smFISH (*n_cells_*, statistical tests and p-value thresholds as in Figure 1B). **D.** Average number of nascent transcripts per cell measured by smFISH (black, error bars: SEM. *n_cells_*, as in Figure 1B-C, KW test – not shown - provides the same pair-wise p-values as in Figure 1C) and normalized nuclear-NF-κB fluorescence intensity. The transcriptional peak is prompt, as it is almost simultaneous to that of NF-κB nuclear localization within our temporal resolution, and sharp, since it is sharper than the peak of NF-κB nuclear localization, as evaluated by linear interpolation as the time t_1/2_ between maximal activation, max(*A*), and 0.5 × max(*A*) (right panel, error bars calculated by computing the minimal and maximal slope of the lines passing through the 20’ and 1h time-points).

Surprisingly, despite the observed heterogeneity in mRNA levels and active TS numbers at single cell level, the population average of the nascent transcriptional dynamics was remarkably similar for all genes, peaking at 20 min post stimulation as measured by either smFISH (**Figure 1D**) or intron-targeted qPCR (**Figure S1D** and **Methods**). Published models for NF-κB mediated gene expression suggest that RNAs are generated proportionally to NF-κB nuclear abundance (Lee et al., 2014; Zambrano et al., 2014b). We tested this notion by comparing nascent transcriptional dynamics with the abundance of nuclear NF-κB –a classical measure of NF-κB activation– obtained by immunofluorescence (see **Methods**) at different time points. Similar to previous reports (Lee et al., 2014), nuclear NF-κB accumulated rapidly and rather homogeneously across the cell population, peaking after 20 minutes and then decreasing in the following three hours (**Figure 1D and S1E**). Surprisingly, following its peak at 20 minutes, average nascent transcription decreased faster than nuclear NF-κB abundance (**Figure 1D**): the time t_1/2_ for the average nascent RNA signal to decrease to half of the peak value is ^~^30 min, whereas it is ^~^100 min for the average NF-κB nuclear localization (**Figure 1D**).

Taken together, our data show that the transcriptional activation of NF-κB target genes is gene- and cell-dependent. However, at population level their nascent transcription is *prompt*, since it peaks synchronously with NF-κB nuclear localization within our temporal resolution, and *sharp*, since it decays faster than NF-κB nuclear localization. We then decided to investigate further how these population-level features emerge from single-cell bursting dynamics using a live-cell reporter for nascent transcription.

### A live-cell reporter of NF-κB-driven nascent transcription recapitulates the dynamics of endogenous genes

To monitor transcription induced by NF-κB in single living cells we used the HeLa 128xMS2 cell line (Tantale et al., 2016) (see **Methods**). Briefly, these cells harbor a single integration of a reporter gene containing 128 intronic repeats of the MS2-stem loop that are bound by a phage coat protein fused to GFP (MCP-GFP), such that bright spot within the nucleus denotes an active TS (**Figure 2A**). The reporter gene is under the control of the HIV-1 LTR, which contains two NF-κB binding sites (Stroud et al., 2009); this compares with the promoters of classic NF-κB targets, which typically harbor from 1 to 5 binding sites (Siggers et al., 2010). TNF-α stimulation induces transcription, as assessed by PCR after 1 hour of stimulation with 10 ng/ml TNF-α (**Figure S2A**). We visualized transcription in our cells using a sensitive widefield microscope (see **Methods**), which allows to visualize both the TS and the single molecules of released transcripts (RNAs, see **Methods** and insets of **Figures 2B-C**). Similar to what observed for *IL6* and *TNF*, we found active TS in only a relatively small fraction of cells (20%) 1 hr after TNF-α induction; an additional 20% of cells displayed mature RNAs but not active TS (**Figures 2C and 2D**). This fractional response was confirmed by smFISH using probes targeting the MS2 RNA (**Figure S2B**) and cannot be ascribed to reporter loss, since active TS were present in 10 out of 10 clonal sub-populations generated (**Figure S2C**). Interestingly, as for the endogenous genes, a fraction of unstimulated cells (5%) also displayed active TSs while 20% displayed only released RNAs (**Figures 2B and 2D**), suggesting previous transcriptional activity potentially due to nonzero nuclear NF-κB basal levels or spontaneous activations, as reported (Zambrano et al., 2014a). Importantly, the population average of MS2 nascent transcriptional dynamics is similar that of the selected endogenous genes, and specifically displays a prompt and sharp response (**Figure S2D**). Hence, our MS2 reporter reproduces both single-cell and population-level features of endogenous NF-κB target genes and thus can be considered a faithful tool to study NF-κB regulated transcription.

**Figure 2.**
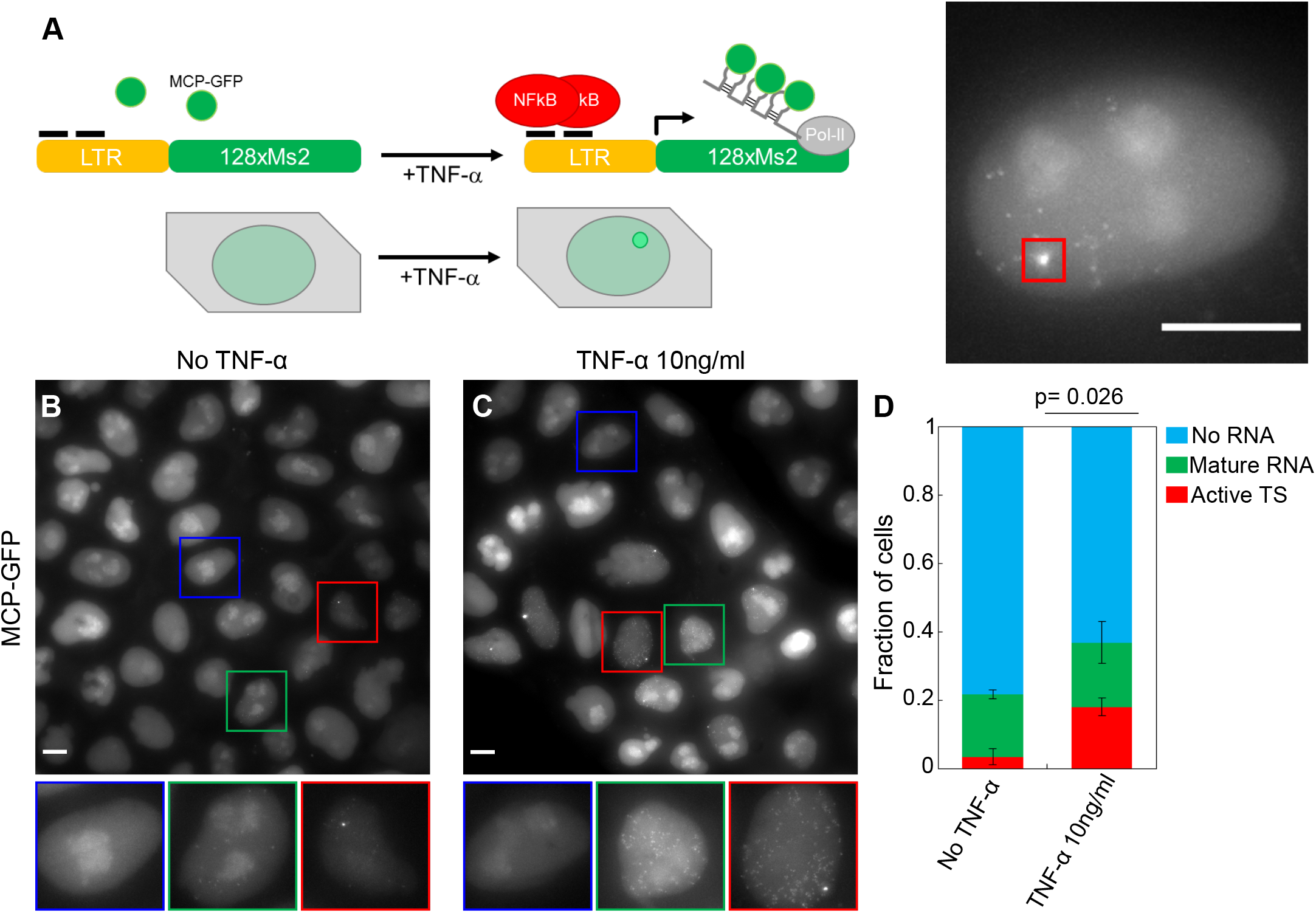
Probing NF-κB transcription in single cells using a MS2 reporter. **A.** The MS2 reporter of TNF-α induced transcriptional activity. 128 MS2 stem loops RNAs are transcribed by the gene under the control of the NF-κB controlled LTR-HIV1 promoter, RNAs are bound by constitutively expressed MCP-GFP protein. As a result, a bright spot appears in the cell nuclei. **B.** Representative image of unstimulated cells observed with a high illumination microscope, allowing to observe cells with a visible active TS (inset, red frame), cells with single RNAs but no visible active TSs (inset, green frame) and none observed (inset, blue frame). **C.** Same for stimulated with 10 ng/ml of TNF-α. **D.** Quantification of the fraction of cells with visible RNAs and visible TSs show statistical difference between TNF treatment or no treatment (mean and SD of 2 independent experiments is plotted, t-test).

### NF-κB mediated transcriptional response is bursty and shaped by a population of “first responders”

We then used our reporter to characterize nascent transcriptional dynamics by monitoring the TS signal in single 128xMS2 cells over time, using a confocal microscope (**Figure 3A**, upper panels). We recorded 3D stacks of 10 μm depth every 3 minutes for 3 hours. A custom software allows to track the cell and detect the TS after a high pass filter of the stack maximal projection (see **Methods** and **Fig. S3_1A**). The maximum signal intensity of the TS is informative of the total TS intensity, since they correlate (see **Figure S3_1B**), while it is independent from the expression level of MCP-GFP in the cell (**Figure S3_1C**). The TS signal is then compared to the MCP-GFP background intensity to distinguish between transcriptionally “active” and “inactive” cells (see **Methods**, **Supplementary Methods** and **Figure S3_1D**). Our time-lapse analyses showed that the MS2 transcriptional activity induced by TNF-α appears as discrete peaks, heterogeneous both in height and frequency, confirming experimentally the “bursty” feature that has been postulated from indirect measurements (Dar et al., 2012; Wong et al., 2018). In addition, “active” and “inactive” cells coexisted both after stimulation **Movie S1 and Fig. 3A**) or no stimulation (**Movie S2 and Fig. 3A**).

**Figure 3.**
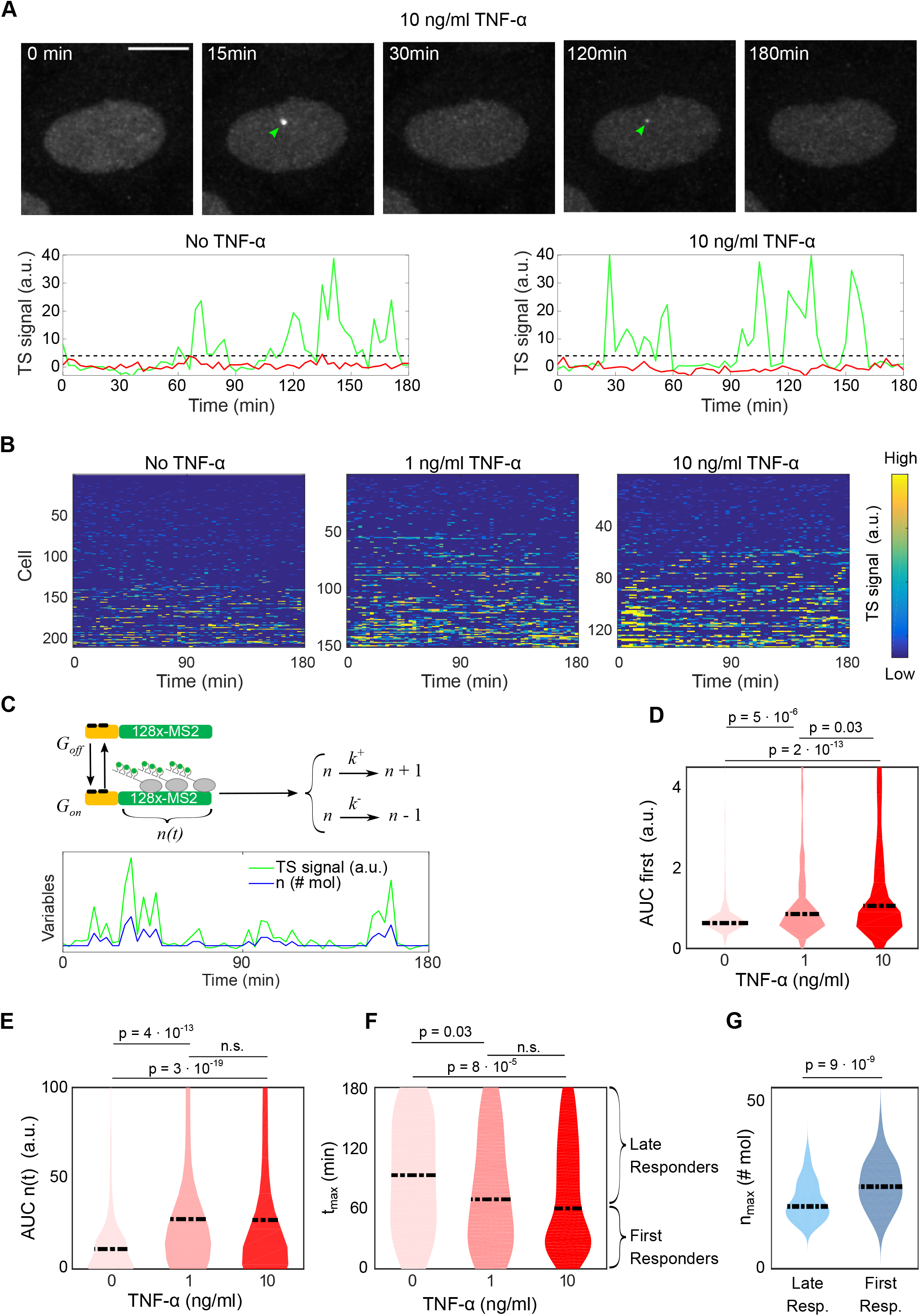
Live cell imaging of MS2 reporter for different doses and stochastic modeling highlights a dose-dependent bursting behavior and the existence of a fraction of first responders to TNF-α. **A.** Exemplary images of a cell stimulated with 10 ng/ml TNF-α and acquired with our live cell imaging setup (maximum projection, scale bar 10 μm). Arrows indicate the detected TS signal. Tracks show TS signal for unstimulated and stimulated cells, either displaying bursts (green) and no bursts (red). Transcribing TS are identified by having signal above or below the threshold (dashed black line) established as four times the standard deviation of the background signals. **B.** TS signal for hundreds of cells, either unstimulated or stimulated with 1 or 10 ng/ml TNF-α, sorted for increasing TS signal. **C.** Scheme of the simple mathematical model of nascent transcription *n(t)* with the activation and inactivation rates of the gene (*k_on_* and *k_off_*), the RNA accumulation rate (*k*^+^) and the RNA release rate (*k*^−^). Example of the inferred transcript levels *n(t)* from a time series of TS signal. **D.** Transcriptional activity during the first burst, and **E**. Transcriptional activity during the whole time course inferred as area under the curve (AUC) of *n(t)*, for the three doses of TNF. **F.** Distribution of the timing of the maximum TS signal, indicating that for 10 ng/ml TNF-α there is a fraction of cells displaying a prompt response. **G.** The peak transcriptional activity *n_max_* is higher in first responders. In all panel, statistical significance is calculated with pairwise Kolmogorov-Smirnov tests.

We repeated the time lapse imaging of our cells for different TNF-α doses and measured TS signals in hundreds of cells (**Figure 3B**). In color-plots, each line corresponds to a single TS observed for 180 minutes and the color reflects the TS signal intensity. The measured transcriptional response is strongly heterogeneous (**Movies S3 to S5**), but controlled by TNF-α, as the timing, the amplitude and the integrated intensity of the detected bursts are modulated by the dose (**Figure S3_2A**), as reported for bulk populations (Tay et al., 2010). Shear stress (Baeriswyl et al., 2019) potentially associated to plain addition of TNF-α-free medium does not lead to observable TS activity (**Figure S3_2B**).

Following previous work, we adapted the random telegraph model of transcription (Suter et al., 2011) to our MS2 reporter gene (**Figure 3C**) (see **Methods** and **Supplementary Methods**) to determine the timespan of gene activations and estimate the evolution of the number of nascent transcripts in time, *n(t)*. The model accounts for the promoter switching between an active and an inactive state with rates *k_on_* and *k_off_*. Once the promoter is in its active state, new transcripts are generated with a rate equal to *k^+^* and processed/realeased with a rate equal to *k^−^*. After verifying that the stochastic model could faithfully infer gene activation from synthetically generated TS time traces (**Figure S3_3A**), we fitted our experimental data with the model (**Figure 3C**), by imposing that the average number of transcripts observed after 20 minutes of stimulation with TNF-α (10 ng/ml) would match with what observed by smFISH (6 RNAs/cell). In agreement with our previous analysis, the amplitude of the first burst is modulated by the dose of TNF-α (**Figure 3D**) and, more generally, the reporter transcriptional activity (estimated as AUC of *n(t)*) increases upon treatment with TNF-α (**Figure 3E**), due to an increase in the gene activation rate *k_on_* and a decrease in the deactivation rate *k_off_* (**Figure S3_3B**).

Importantly, we found a fraction of cells responding almost synchronously and within few minutes after TNF-α stimulation; this first response occurred earlier upon higher TNF-α (**Figure 3F**). “First responders”, which can be defined as those cells reaching a maximum transcriptional activity before the median value (60 mins for 10 ng/ml TNF-α), have a stronger transcriptional activity than the other cells (**Figure 3G**). After this first burst of transcription, intrinsic stochasticity dominates the individual cell response, as can be quantified by the evolution in time of the coefficient of variation of the number of nascent transcripts *n(t)*. The coefficient of variation has a minimum at 20 minutes (**Figure S3_3C**), which indicates an early synchronous round of transcription in a fraction of cells. The combination of the transcriptional activity of these first responders and the increased synchronicity of bursting at approximately 20 min post TNF-α lead to the observed prompt transcriptional response at population level.

### First responders are not purely stochastic

We next used our live-cell reporter to characterize to what extent the responses to TNF-α were actually stochastic. Using our previously described microfluidics setup (Zambrano et al., 2016), we challenged our MS2×128 cells with two independent 1 hour pulses of 10 ng/ml TNF-α separated by a 2 hours washout (see **Methods**) and followed TS activity in hundreds of cells (**Fig. 4A**). Similarly to what we observed for a single stimulation, the bursting parameters extracted from this two-pulses experiment were found to be modulated by TNF-α (**Figure S4A**). We then determined the fraction of cells responding to the first, to the second, and to both pulses (**Figure 4B** and **Movie S6**). A majority of responding cells responded to both pulses, and a fraction of cells responded to only one. Surprisingly, the fraction of cells responding to both pulses is significantly higher than what could be expected from statistically independent transcriptional activations (**Figure 4B** and **Supplementary Methods**). Moreover, the maximum TS signal, expressed as number of nascent transcripts *n_max_* for each pulse, was higher for cells responding to both TNF-α pulses than for cells responding to only either one of them (**Figure 4C**); the AUC behaves analogously (**Figure S4_B**). Further, the timing to the maximum TS signal (*t_max_*) after a TNF-α pulse was shorter on average for cells that respond to both pulses than for cells that respond to just one (**Figure 4D**), and similar to the *t_max_* of the previously identified “first responders”.

**Figure 4.**
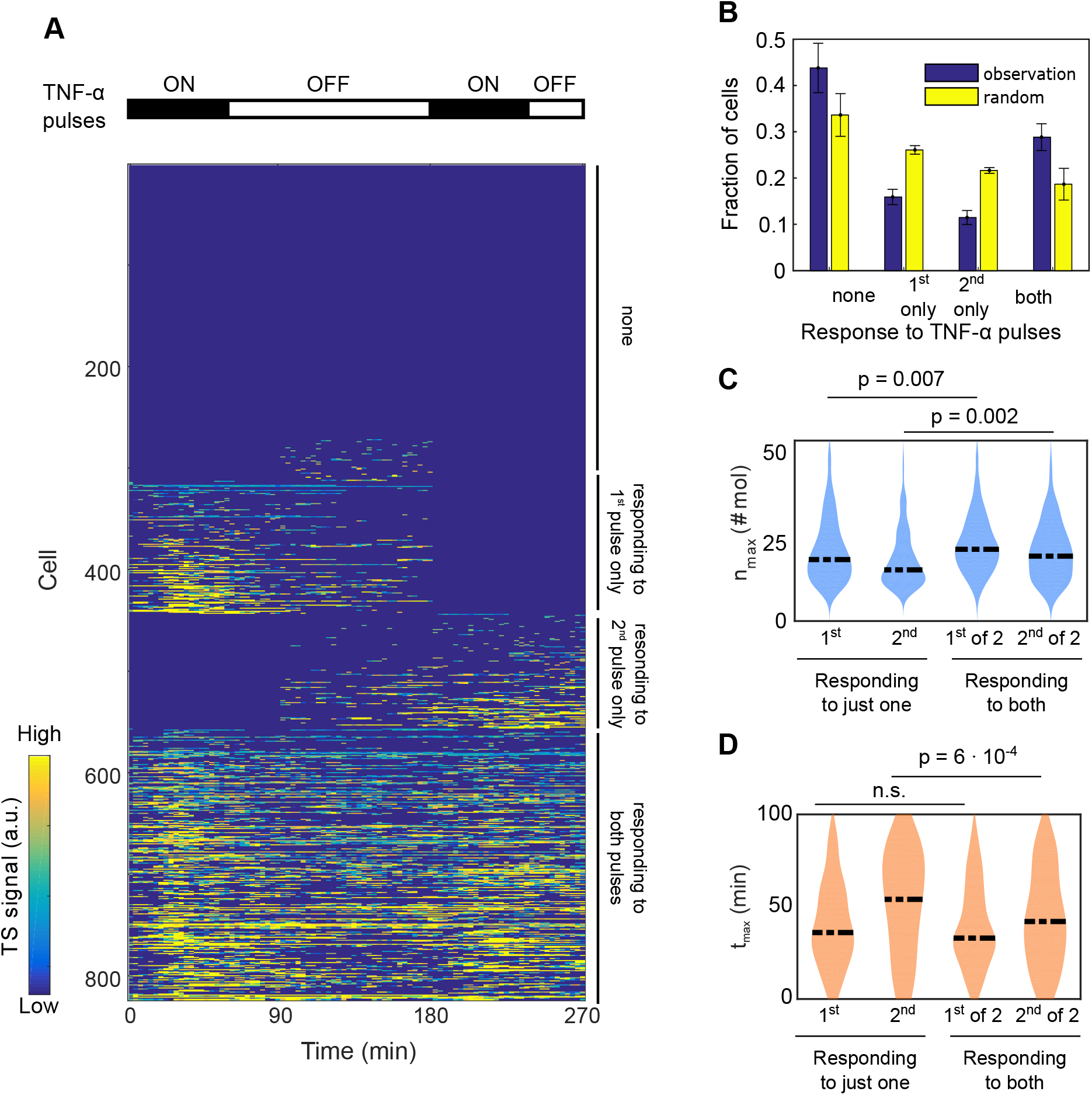
Pulsed TNF-α stimulation shows that transcriptional bursts are not purely stochastic. **A.** TS signal for hundreds of cells for after two pulses of one hour of 10 ng/ml TNF-α separated by a two hours washout, sorted for increasing TS signal. Cells are clustered as non-responding – within 90 minutes of each pulse–responding to only one of the two pulses or responding to both pulses. **B.** Fraction of cells responding to none of the TNF-α pulses, just the first or just the second (mean and standard deviation of 3 independent experiments), and predicted fraction for statistically independent activation (random). **C.** Maximum TS signal (in number of transcripts) after the first and second pulse for the sub-populations identified above. Cells responding to both TNF–α pulses display a stronger response to both the first and the second pulse. **D.** The timing of the maximum of the TS signal after each TNF-α pulse, indicating that cells that are primed to response do so more quickly upon the first pulse that the remaining populations, in particular those responding only to the first or the second.

Overall, our results suggest that despite an intrinsic stochasticity in gene activation (cells can respond to either one TNF-α pulse or to both) there is a higher than expected proportion of cells that respond to both pulses, which excludes the statistical independence of the two responses. The data indicate that some cells are in a “first responder” state lasting longer than 180 minutes; first responders are activated faster, higher and more often than other cells.

### The timing of the nascent transcriptional response does not depend on NF-κB nuclear localization dynamics

Once we established that the population-level transcriptional response to TNF-α is the result of heterogeneous transcriptional activation in single cells, we asked whether the latter emerged from heterogeneous nuclear localization dynamics of NF-κB. We stably transfected our MS2×128 cells with a previously validated RFP-p65 construct (Bosisio et al., 2006) (**Figure 5A**) and measured concomitantly TS signal intensity and NF-κB nuclear localization (see **Methods**) in single living cells (see **Figure 5A** and **MovieS7**). NF-κB localization dynamics were similar for all the responding cells (see **Methods**), as previously reported (Lee et al., 2014; Tay et al., 2010), whereas the transcriptional response was heterogeneous, as shown in previous experiments (**Movies S8-S10**). Parameters governing the bursting kinetics are similar to those obtained from untransfected cells (**Figure S5A**), excluding an effect of transfection on results. At the single cell level, the change in NF-κB nuclear abundance does not correlate with the amount of nascent transcription (r^2^=0.0055) (**Figure S5B**). Thus, uniform nuclear translocation of NF-κB drives highly non-uniform transcription at single cell level, which highlights the stochastic nature of the transcriptional activation process.

**Figure 5.**
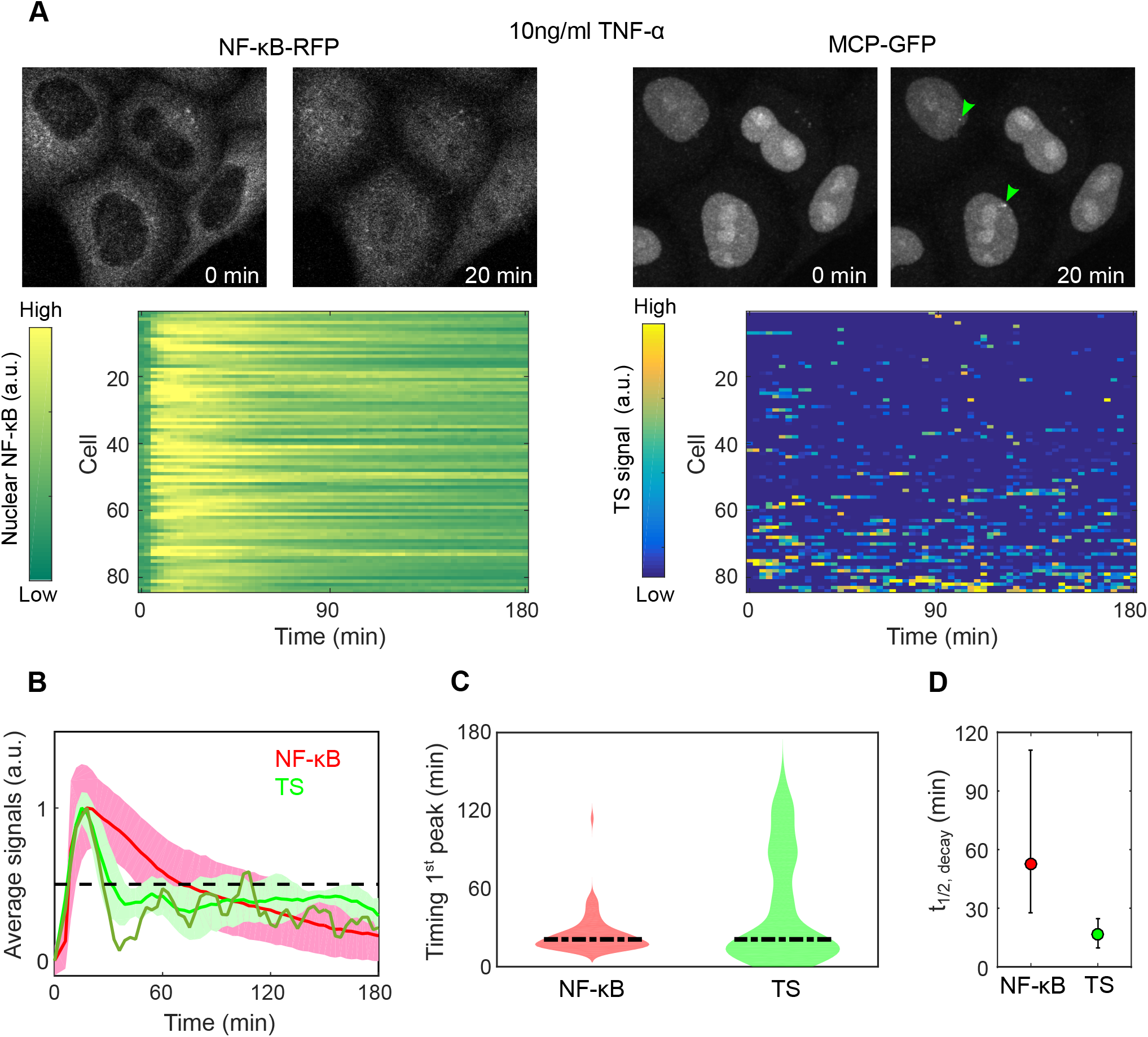
Simultaneous imaging of NF-κB translocation and MS2 transcriptional activity highlights the promptness and sharpness of the transcriptional response. **A.** Top: exemplary images of cells stimulated with 10 ng/ml TNF-α before and after 30 minutes stimulation with 10 ng/ml TNF-α. Note the activation of NF-κB in all the cells, while the TS appears active in the indicated ones (arrows) at that specific time point. Bottom: TS signal activity and nuclear NF-κB activation for hundreds of cells sorted for increasing TS signal. **B.** Plot of the normalized average TS signal for three experiments of (green, standard deviation is represented), superimposed with the average NF-κB nuclear intensity assessed by live cell imaging (red) with standard deviation inferred from imaging data. The dark green line represents the TS activity of transfected cells, within the range of variability observed for untransfected cells. The plot indicates that both signals peak simultaneously but TS activation decrease more sharply. **C.** The quantification of the timing of the first maximum of NF-kB nuclear localization and of TS activity, showing that the medians are similar but the latter is more heterogeneous with prompt and late responders. **D.** Estimation of the variability of the decay time t_1/2_ for the TS signal and NF-κB nuclear localization, obtained from panel B. The decay time of the transcription signal is much lower than that of NF-κB, indicating a sharper response.

Such time-resolved measurements allowed us to quantify more finely the promptness of the transcriptional response. By superimposing the average data for NF-κB translocation and MS2 reporter transcriptional activity we found that both peak almost simultaneously at about 20 minutes post stimulation (both for transfected and untransfected cells, **Figure 5B).** The timing of the first NF-κB nuclear translocation peak (typically the only one, see **Figure S5C**) matched with the timing of the first peak of nascent transcription (**Figure 5C**) for most of the cells previously identified as “first responders”(**Figure 5C**). Nuclear NF-κB is therefore the limiting factor for transcriptional activation. This is compatible with the observation that NF-κB can find its targets rapidly (search time ^~^2 min), as can be derived from recent single molecule imaging data(Callegari et al., 2019) (see **Supplementary Methods**). We also quantified the sharpness of the nascent transcriptional response and of NF-κB localization by computing their time t_1/2_. The TS signal decayed faster than NF-κB nuclear abundance, in agreement with what observed for endogenous genes by smFISH. Thus, sharpness is reproduced faithfully by time lapse imaging of our reporter gene (**Figure 5D**).

In short, these results illustrate how the nascent transcriptional response to TNF-α is more heterogeneous than NF-κB nuclear localization among the cells in the population. Moreover, a fraction of prompt-responder cells is responsible for the prompt and sharp transcriptional response emerging at population level.

### A model combining NF-κB mediated gene activation and a refractory state recapitulates the prompt and sharp nascent transcriptional response

To gain insights on the origin of the prompt and sharp NF-κB mediated transcriptional response, we explored mathematical models for NF-κB-driven transcription. We performed stochastic and deterministic simulations of gene activity (see **Supplementary Methods**) and compared their results to our experimental data. A first candidate for our exploration was the random telegraph model of transcription, where the gene switches between on and off states in a purely stochastic fashion, with constant switching rates. This model however could not recapitulate our experimental data. For example, the experimentally measured gene off-times are described by a unimodal distribution with a shape that varies between unstimulated and stimulated conditions (**Figure 6A**), rather than by the exponential that would be expected from the random telegraph model (Model 0, **Figure 6B**). Two alternative mechanism have been suggested to give rise to these unimodal distributions: a) the presence of a gene refractory state (Molina et al., 2013; Suter et al., 2011) that prevents the gene from immediately starting a second round of transcription after the first one is over and b) an oscillatory transcription activator-dependent modulation of the gene activation (Zambrano et al., 2015). As shown below, none of these two models can independently reproduce the experimental features that we observed, but their combination can.

**Figure 6.**
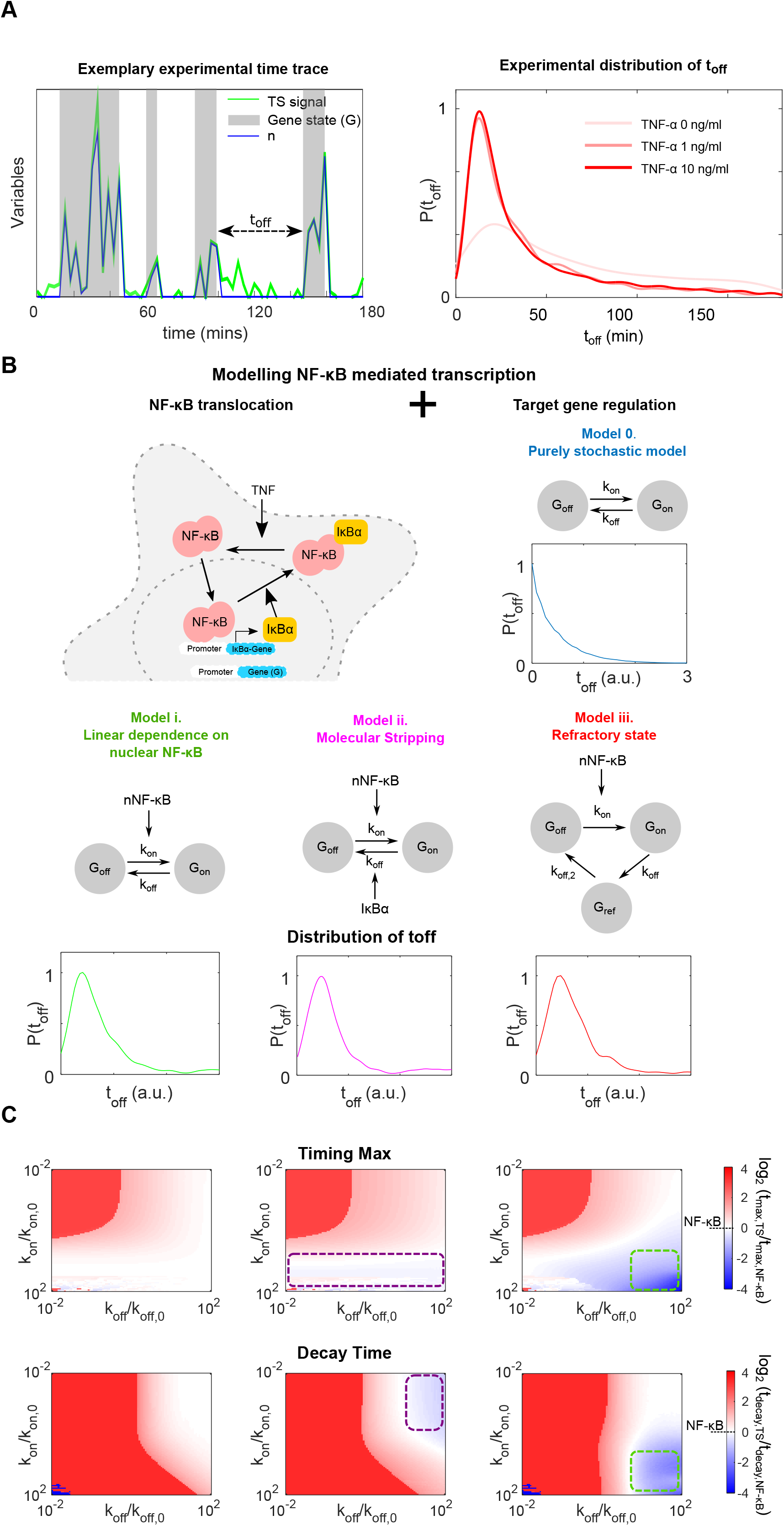
Identification of a minimal mathematical model recapitulating NF-κB mediated transcription dynamics. **A.** Example of the inferred transcript levels *n(t)* given a TS signal time series. The off times t_off_ are computed as described (top). Unimodal distribution of the off times obtained from our experimental data (bottom). **B.** Scheme of a simple mathematical model where gene activation is modulated by NF-κB while inactivation is governed by the concentration of the inhibitor IκBα. Different possible mechanisms of activation of the target gene are considered. The classical telegraph model of transcription (Model 0) with constant activation and inactivation rates gives rise to exponential distribution of the off times (inset) so cannot describe experimental data. Based on the literature and our observation we propose alternative models: linear activation (Model i), molecular stripping (Model ii) and gene with a refractory state (Model iii). All of them reproduce the unimodal distribution of *t_off_* (insets). **C.** We screened the timing of the peak of the gene activity (top panels) and the sharpness (bottom panels) of the peak for Models i to iii, two orders of magnitude above and below reference values (k_on,0_ and k_off,0_). The color-code indicates the promptness and the sharpness of the peak, respectively, as compared to the peak of nuclear NF-κB. Model i does not give prompt and sharp responses. Model ii gives prompt responses in a region that does not overlap with the region giving a sharp response, both highlighted with a purple square. Finally, Model iii with NF-κB mediated activation and a refractory state is the only one giving parameters combination (high k_on_ and k_off_) leading to a prompt and sharp transcriptional response, highlighted with a green square.

In previous explorations we simulated NF-κB response to TNF-α using a simple mathematical model (Zambrano et al., 2014b) (see **Figure 6B** and **Supplementary Methods**); here, we analyzed the transcriptional dynamics of a prototypical target gene by modelling different NF-κB controlled gene activation-deactivation schemes inspired by experimental observations, among which our own. We used deterministic modeling to simulate population-average gene activity dynamics (**Figure S6A**) and stochastic modeling to simulate bursty stochastic transcription at single cell level, including the distribution of the off times (**Figure S6B**). The key parameters considered are the gene inactivation (*k_off_*) and activation rates (*k_on_*), which we varied four orders of magnitude around values used in the literature (Tay et al., 2010; Zambrano et al., 2014b) (see **Supplementary Methods**). To constrain our exploration, we modeled first the gene activation rate as depending linearly (non-cooperatively) on NF-κB nuclear concentration, as proposed in a number of models (Tay et al., 2010; Zambrano et al., 2014b) and deduced from previous experiments and thermodynamic considerations(Siggers et al., 2010) (Model i, **Figure 6B**). This model allows to reproduce the non-monotonocity of the off times observed experimentally (**Figure 6B**), as predicted (Zambrano et al., 2015), but is unable to reproduce the prompt and sharp gene activation observed in our experiments (**Figure 6C**).

A recently proposed mechanism that in principle could rapidly shut down transcriptional activity and produce “sharpness” is molecular stripping, by which IκBα actively induces the dissociation of NF-κB from its binding sites on DNA (Dembinski et al., 2017; Potoyan et al., 2016) (Model ii, **Figure 6B**). A model based on molecular stripping reproduces the unimodal distribution of the inactivation times (**Figure 6B**) and we could indeed identify a sector of parameter space –low *k_on_* and high *k_off_* values– resulting in sharp transcriptional responses at the population level (**Figure 6B**). However, these parameters were not compatible with a prompt transcriptional activation, which was found for high *k_on_* values instead (see purple areas in **Figure 6C** and examples in **Figure S6C**). To test these predictions, we co-treated our cells with TNF-α and cycloheximide (CHX), which blocks protein synthesis and hence IκBα synthesis and stripping. CHX is effective as demonstrated by the progressive decay observed in the nuclear fluorescence of MCP-GFP (**Movie S11**) and by higher NF-κB nuclear localization post TNF-stimulation (**Figure S6D**), as expected from blocking IκB re-synthesis. However, the TS signal decay after reaching its maximum *t_max_* remains almost unchanged by CHX, indicating that it does not depend on IκBα re-synthesis and stripping (**Figure S6E**).

Finally, we tested a model that combines the two previously mentioned mechanisms: NF-κB mediated activation by nuclear translocation and a gene refractory state (Model iii, **Figure 6B**). As previously reported(Molina et al., 2013), such model reproduces the non-monotonous distribution of “off times” of our bursty transcription data (**Figure 6B**). Interestingly, we find a wide region of parameter space (characterized by high *k_on_* and *k_off_*) compatible with both prompt and sharp gene activation (see green squared areas in **Figure 6C** and examples in **Figure S6F**). Furthermore, the simulated bursts have a structure clearly reminiscent of our experimental data, differently from the ones obtained from the other models (**Figure S6G**). Importantly, such model is able to reproduce two key features: (i) the temporal evolution of the coefficient of variation of nascent transcription that we observed experimentally, with maximum synchronization of the bursts approximately 20 min post-stimulation (**Figure S6H**), and (ii) the presence of a fraction of first responders in the cell population (**Figure S6I)**. When using the experimentally determined NF-κB nuclear dynamics as input to simulate the gene activation rates of each single cell following the scheme of Model iii, we also reproduced a population-level prompt and sharp nascent transcriptional response (**Supplementary methods** and **Figure S6J**).

Hence, a refractory gene state is necessary to recapitulate the experimentally determined features of NF-κB mediated nascent transcription upon TNF-α, including a prompt and sharp transcriptional response emerging from a fraction of first responders.

## Discussion

NF-κB dynamics is fundamental for the proper temporal development of inflammation. Previous reports (Ashall et al., 2009; Nelson et al., 2004; Sung et al., 2009) had shown that the NF-κB mediated transcriptional response to TNF-α can display a variety of dynamics, including genes whose mature transcripts peak early (at 30 min) or late (>3 hours), and even oscillating and non-oscillating gene expression patterns(Zambrano et al., 2016). We and others (Hao and Baltimore, 2013; Zambrano et al., 2016) suggested that such mRNA expression patterns arise from a common nascent transcriptional response, that peaks typically 20-30 minutes post stimulation. However, all these observations were based on population-level transcriptional measures, so how single-cell transcriptional response contributes to these features remained an open question that we have addressed in this work.

### Different endogenous genes are expressed with different degrees of variability among individual cells upon TNF-α, but share a common population-level prompt and sharp nascent transcriptional response

Using single-cell smRNA-FISH for three bona-fide NF-κB target genes at different time points post TNF-α stimulation, we found that all of them were expressed heterogeneously across the population, although *NFKBIA* (coding for the inhibitor IκBα) was expressed more uniformly than *IL6* and *TNF*, coding for cytokines. Surprisingly, though, we found that the population dynamics of the nascent transcriptional response was very similar for these three genes, in spite of their marked differences in expression level and variability at single cell level, with Gini coefficients ranging between 0.2 to 0.5. Concomitant measurement of NF-κB nuclear localization by immunofluorescence showed that such common nascent transcriptional response is *prompt*, peaking simultaneously to NF-κB nuclear abundance, and *sharp*, decaying faster than the peak of NF-κB nuclear localization.

### Population-level promptness and sharpness arises from heterogeneous bursting in single cells, including a fraction of “first responders”

NF-κB response to TNF-α has been described as digital, giving rise to a transcriptional output proportional to the fraction of responding cells (Tay et al., 2010), which suggested a relatively uniform transcriptional response across the population. Instead, using our MS2 nascent transcription reporter we find that a digital activation of NF-κB in our cells (100% responding to 10 ng/ml of TNF-α, assessed by immunofluorescence and live cell imaging) gives rise to an extremely heterogeneous transcriptional response. This includes a fraction of “first responders”, cells that reach a maximum transcriptional response higher and earlier than the other cells, and are more likely to respond to consecutive pulses of TNF-α. Interestingly, a fraction of “first responders” was identified when studying cellular responses to viral-activated interferon-beta signaling (Patil et al., 2015). sm-FISH data for endogenous genes *NFKBIA*, *IL6* and *TNF* also confirm a peak of TS activity for a fraction of cells within 20 minutes. We ascribe the nascent transcriptional response observed at the population level to first responders that start transcribing earlier and more strongly than the other cells. Such rapid surge in nascent transcription is compatible with a short NF-κB search time on chromatin. Hence transcriptional initiation can indeed occur nearly simultaneously to NF-κB translocation in the nucleus, as we observe experimentally in some cells.

### The transcriptional response to TNF-α has a stochastic component that is relevant to HIV latency

When challenging our cells with two pulses of TNF-α we find that while some cells respond to both pulses, some will respond just to the first or the second. Our cells harbor an LTR-HIV1 promoter, therefore this observation could represent the microscopic equivalent of a recently identified mechanism involved in HIV1 latency, by which proviruses not induced after a first stimulation can be induced after by a second one (Ho et al., 2013). This mechanism leads to a stochastic latency exit and it is clinically important as it may prevent curing patients from the virus by the “shock-and-kill” approach.

### Analysis of transcriptional bursts highlights the existence of a characteristic inactive time after each gene activation

Our live cell imaging analysis of nascent transcription shows that after gene activations –during which multiple burst of transcription can occur– there is typically a gene inactive time of approximately 25 minutes. This is characterized by a unimodal distribution of the gene “off” times obtained from our stochastic inference framework. Our previous theoretical work (Zambrano et al., 2015) shows that such characteristic unimodal distribution can in principle arise from NF-κB-driven gene activation in a gene that has just two states (2-states model). The same distributions were observed by others (Molina et al., 2013; Suter et al., 2011; Tantale et al., 2016) and modelled by adding an additional gene refractory state (3-states model). A study of our gene reporter under the control of HIV TAT protein suggested that a non-permissive state on the timescale of tens of minutes can be related to the dissociation of TBP from the promoter (Tantale et al., 2016). However, neither the 2-state model (where inactivation is either spontaneous or driven by the inhibitor IκBα through “molecular stripping” (Potoyan et al., 2016)) nor the 3-state model (including a refractory state) can reproduce by themselves our key experimental findings of promptness and sharpness.

### Only a mathematical model combining both NF-κB driven gene activation and a refractory state can reproduce experimental observations of promptness and sharpness

Instead, by combining NF-κB mediated activation and a gene refractory state, the experimentally observed dynamics of transcription are reproduced. This model also reproduces other features in our experiments that two-state models cannot, such as the existence of “first responders” and a peak of bursting synchrony at 20 minutes post-stimulus. Overall, our model illustrates how a simple 3-state dynamics can produce a heterogeneous transcription activity at single-cell level and at the same time a sharp population-level transcriptional output.

### Sharp and prompt nascent transcriptional responses emerging from a fraction of “first responders”: a general feature for inducible transcription factors?

Previous population-level work on transcription suggested that gene-specific NF-κB driven expression profiles are mostly controlled by mRNA processing and degradation (Hao and Baltimore, 2013, 2009), while nascent transcription dynamics are shared among the different genes (Zambrano et al., 2016). Our work reinforces this viewpoint with a single-cell perspective, since we show how such similar nascent transcriptional dynamics emerge from prompt and bursty transcription in single cells. If mRNA degradation controls the temporal evolution of gene expression, a prompt and sharp peak of nascent transcription is a better-suited input to generate gene-dependent expression profiles as compared to a slowly varying transcriptional activity. The observed refractory state might have evolved from the necessity of sharpening the inherently stochastic transcriptional process, providing an opportunity window of decision (Zambrano et al., 2016). Furthermore, it is enough to provide a fraction of “first responders”, which might be useful to temporally stratify the population response to stimuli.

Other inducible transcription factors such as p53 have similar search times (Loffreda et al., 2017) to the one we calculated for NF-κB and produce population-level gene-independent nascent transcription dynamics and gene-specific mRNA profiles due to differential RNA degradation (Hafner et al., 2017; Koh et al., 2019; Porter et al., 2016). It is then tempting to speculate that other transcription factors that need to respond rapidly to intracellular (e.g. p53) (Hafner et al., 2017) or extracellular cues (e.g. STAT3, GR)’(Alonzi et al., 2001; Stavreva et al., 2019) might exploit a similar design principle to produce a time-resolved, prompt and sharp nascent transcriptional response.

In conclusion, our data and models show how the expression of NF-κB target genes can be coordinated at cell population level and yet be heterogeneous across single cells, and further provide a framework for understanding how transcription factors can achieve prompt and sharp transcriptional responses.

## Acknowledgements

The research leading to these results has received funding from Fondazione Cariplo (A.L. and D.M.: 2014-1157 to D.M.), OSR (OSR Seed Grants S.Z. and D.M.) and the Italian Cancer Research Association (AIRC, D.M. IG 2018-21897, A.A., F.C., E.C. and S.Z. IG 2017-18687 to A. A.).

## Author contribution

Conceptualization: SZ, NM, DM

Supervision: SZ, AA, MEB, CT, DM

Visualization: SZ, AL, NM, DM

Formal Analysis: SZ, AL, AA, MEB, NM, DM

Investigation: SZ, AL, EC, GS, FC, EB, NM, DM

Writing-Original draft: SZ, DM

Writing-review and Editing: SZ, EB, CT, AA, MEB, NM, DM

Funding acquisition: SZ, CT, EB, AA, MEB, DM

## Declaration of Interests

The authors declare no competing interests

## STAR Methods

### Key Resources Table

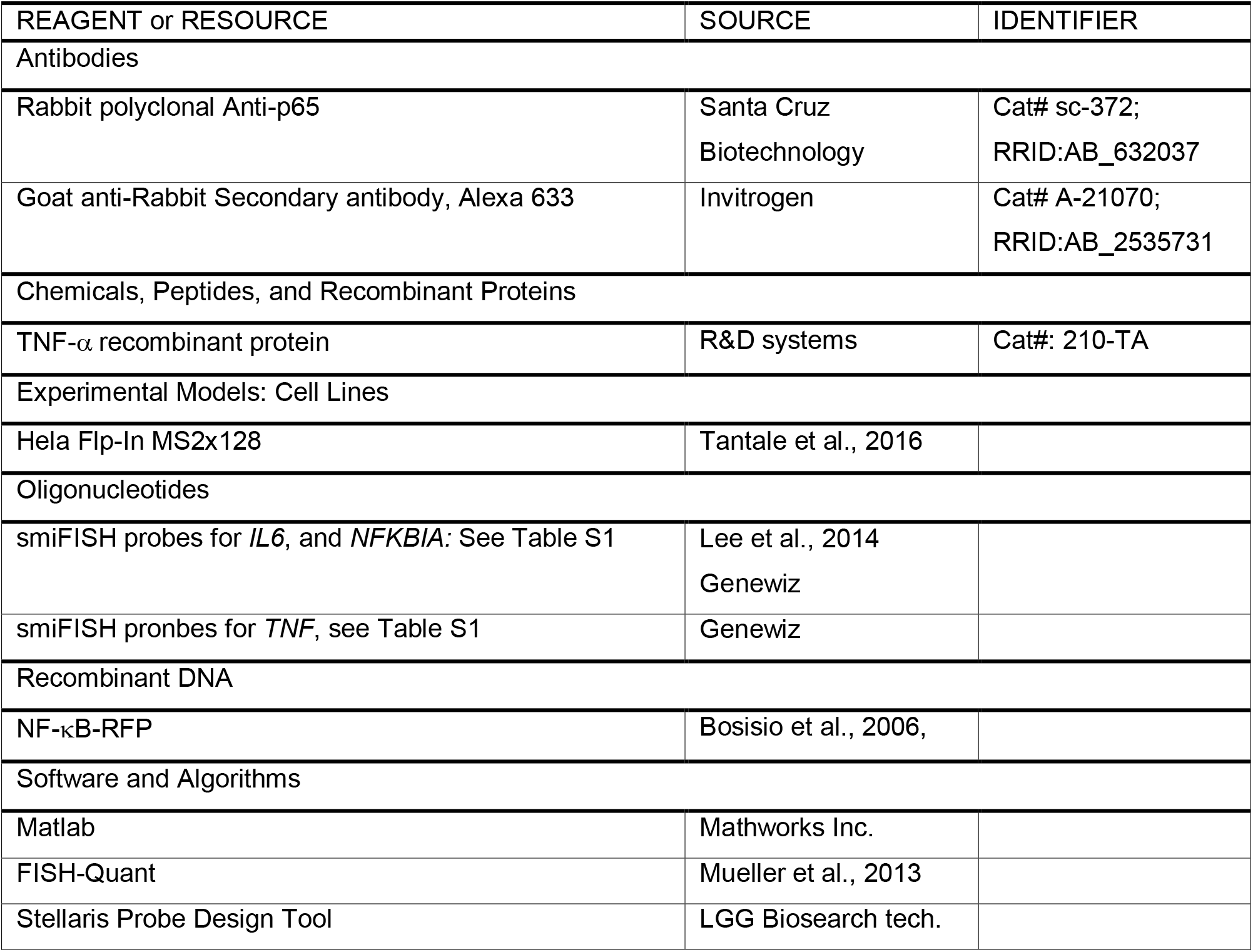

| REAGENT or RESOURCE | SOURCE | IDENTIFIER |
| --- | --- | --- |
| Antibodies |  |  |
| Rabbit polyclonal Anti-p65 | Santa Cruz Biotechnology | Cat# sc-372;<br>RRID:AB_632037 |
| Goat anti-Rabbit Secondary antibody, Alexa 633 | Invitrogen | Cat# A-21070;<br>RRID:AB_2535731 |
| Chemicals, Peptides, and Recombinant Proteins |  |  |
| TNF-α recombinant protein | R&D systems | Cat#: 210-TA |
| Experimental Models: Cell Lines |  |  |
| Hela Flp-In MS2x128 | Tantale et al., 2016 |  |
| Oligonucleotides |  |  |
| smiFISH probes for *IL6*, and *NFKBIA*: See Table S1 | Lee et al., 2014<br>Genewiz |  |
| smiFISH probes for *TNF*, see Table S1 | Genewiz |  |
| Recombinant DNA |  |  |
| NF-κB-RFP | Bosisio et al., 2006, | |
| Software and Algorithms |  |  |
| Matlab | Mathworks Inc. |  |
| FISH-Quant | Mueller et al., 2013 |  |
| Stellaris Probe Design Tool | LGG Biosearch tech. |  |

### Resource Availability

#### Lead Contact

The lead contact for this study is Samuel Zambrano.

#### Materials Availability

All unique reagents generated in this study are available from the corresponding authors with a completed Material Transfer Agreement.

#### Data and Code Availability

The stochastic simulation and inference software is available at: https://github.com/MolinaLab-IGBMC/ All the software generated is provided upon request.

### Experimental Model and Subject Details

A clonal population of HeLa Flp-in H9 cells constitutively expressing MCP-GFP and with a single integration of the HIV-1 reporter gene was created using the Flip-In system (Life Technologies, Carlsbad, USA) as previously described (Tantale et al., 2016). Cells were cultured in phenol-red free DMEM, supplemented with 10% FCS, 1% L-Glutamine and Pen/Strep at 37°C with humidified 5% CO2. Hygromycin (150 μg/ml, Sigma Aldrich, St. Louis, USA) to guarantee the continuity of Flp-In integrants. The isolation of individual clones has been obtained by limiting dilution in a 96 well plate. Cells stably expressiong RFP-p65 were generated after transfection with the plasmid described in (Bosisio et al., 2006), antibiotic selection and sorting.

TNF-α: Mouse recombinant TNF-α (R&D Systems, Minneapolis, USA) was diluted in cell culture medium prior to the injection. CHX was used at 5mg/ml and diluted in cell culture medium concomitantly to TNF-α.

### Methods Details

#### Immunofluorescence

HeLa 128xMS2 plated on glass coverslips were induced with 10 ng/ml of TNF-α and where specified treated with CHX. At the indicated time points coverslips were fixed in 4% paraformaldehyde for 10 min a room temperature (RT), washed with 150mM of NH_4_Cl for 15 min and permeabilized with 0.1% Triton X-100. Coverslips were blocked in PBS 5% BSA 20% FBS for 1 hour at RT and probed with a p65 antibody (SC-372, Santa-Cruz Biotechnology Inc, Dallas, USA) diluted 1:200 at 4 °C overnight. Coverslips were washed three times in washing buffer (PBS 0,2% BSA 0,05% Tween-20) and incubated for 1 h a RT with the secondary anti-rabbit antibody AlexaFluor 633 (Life Technologies) diluted 1:1000. All the antibodies were diluted in washing buffer. Following DNA staining with 1 μg ml^−1^ Hoechst 33342 (Hoechst AG, Frankfurt, Germany) in PBS, the coverslips were mounted on glass slides using Vectashield (Vector Laboratories, Peterborough, UK) mounting media. Nuclear concentration was quantified for hundreds of cells from these images using previously described MATLAB routines (Zambrano et al., 2014a), available upon request.

#### PCR

For RNA isolation cell culture samples were collected in Trizol (Invitrogen) and purified using Nucleospin RNA kit (Macherey Nagel, Duren, Germany). RNA quantity and purity was checked using a NanoDrop fluorometer (Thermo Fisher Scientific, Walrtham, USA) and to control the integrity of total DNA an aliquot of the samples was run on a denaturing agarose gel stained with SYBR Safe (Thermo Fisher Scientific). To assess co-transcriptional or post-transcriptional splicing, RT was performed with either oligo-dT or with random primers using High Capacity cDNA Reverse transcription kit (Thermo Fisher Scientific). Competitive 3-primers PCR was performed as described in (Tantale et al., 2016).

#### qPCR

RNA was extracted and quantified as described above. Reverse transcription of 2 μg of RNA was performed according to the manufacturer’s instructions using the QuantiTect reverse transcription kit (Qiagen, Hilden, Germany). The PCR reaction were done in LightCycler 480 SYBR Green I Master mix (Roche, Basel, Switzerland). Melting curve analyses were carried out to ensure product specificity, and data were analyzed using the 2^−ΔΔCt^ method. Relative nascent RNA expression levels were normalized to *glyceraldehyde3’phosphate dehydrogenase* (*GAPDH*). The primers used for qPCR were: NFKBIA-FW: 5’-ACCTGGCCTTCCTCAACTTC-3’; NFKBIA-REV: 5’-AGGATGTGGGCTGATGTGAA-3’; IL6-FW: TGTGAAAGCAGCAAAGAGGC-3’; IL6-REV: 5’-TGCATGCAAGAGGGAGAAGT-3’; MS2-unspliced-FW: 5’-AATGGGCAAGTTTGTGGAATTGGTT-3’; MS-2-spliced-FW: 5’-CGAACAGGGACTTGAAAGCGA-3’; MS2-REV: 5’-GATACCGTCGAGATCCGTTCA-3’.

#### smFISH

In-situ hybridization was carried out according to the smiFISH (single molecule inexpensive FISH) approach where unlabeled primary probes are prehybridized to a secondary common fluorescently labelled probe (Tsanov et al., 2016). Primary smiFISH probes for *NFKBIA* and *IL6* were designed as in (Lee et al., 2014). smiFISH probes for *TNF*, were designed with the Stellaris FISH online tool (LGC Biosearch Technologies, Oddeson, UK). A FLAP sequence was appended at the 3’ of each probe. The complementary FLAP probes, labelled with Cy5, were hybridized to primary probes as described in (Tsanov et al., 2016). Cells were fixed in 4% PFA for 10 min at RT, then washed twice in PBS and permeabilized in cold 70% EtOH at −20 °C overnight. The day after, coverslips were washed twice with washing buffer I (10% Saline Sodium Citrate (SSC) in RNase-free water) and once in washing buffer II (10% SSC, 20% formamide solution, diluted in RNase-free water). Cells were incubated overnight with the hybridized flap-structured duplex in a humidified chamber at 37 °C. The probes were diluted 1:100 in the hybridization buffer (10% (w/v) of dextran sulfate, 10% of SSC-20× buffer and 20% formamide in Rnase-free water). Following the hybridization, cells were washed twice in buffer II in the dark for 30 min at 37 °C, then washed in PBS for 5 min and stained with 1 μg/ml Hoechst 33342 in PBS. The coverslips were then mounted on glass slides using Vectashield (Vector Laboratories, Peterborough, UK) mounting media. The sequences of the probes used for smiFISH are provided in Supplementary Table 1. For the MS2 transcripts only, smFISH was carried out using the protocol and the probes described in (Tantale et al., 2016).

Imaging was performed on a custom-built widefield microscope with single molecule sensitivity, by using a led source for illumination (Excelitas Xcite XLED1, Qioptiq, Rhyl, UK), a 60x 1.49NA Olympus objective (Olympus Life Science, Segrate, IT), and an Hamamatsu Orca Fusion sCMOS detector (Hamamatsu Photonics Italia S.r.l, Arese, Italy) resulting in a pixel size equal to 108nm. For every field a z-stack series of images were acquired with 0.3 μm step size, to count the number of mature RNAs for each cell.

#### Live cell imaging

*<u>Widefield Microscopy</u>. 3D Stacks were collected using the microscope described above, using a step-size of 0.3* μm, a 100x 1.49 NA objective and a Photometrics Evolve EM-CCD camera (Teledyne Technologies Inc., Thousand Oaks, USA) resulting in a pixel size of 158 nm. The microscope was equipped with a temperature and CO_2_ control for this purpose (Okolab, Naples, Italy)._ <u>*Confocal microscopy:*</u> 16 bit-1024×1024 pixels images were acquired using 63X objective on a Leica SP5 (Leica Microsystems, Wetzlar, Germany) confocal microscopy with temperature and CO_2_ control, as described elsewhere (Sung et al., 2009). For each time point of our time-lapse (sampled every 3 minutes) and for each position of the sample considered we collected z-stacks composed by up to 16 slices, with a step of 0.7 μm, to acquire the whole thickness of the sample.

#### Microfluidics

We used the CellASIC^®^ ONIX Microfluidic Platform as described previously (Zambrano et al., 2016): cells were plated one day before the experiment in CellASIC™ ONIX M04S-03 Microfluidic Plates, consisting of microfluidic wells connected through channels to a series of reservoirs (inlets) containing media with selected concentrations of TNF-α stimuli that can be flown through the chambers. To avoid cell stress or toxicity, the microfluidic plates are primed with 10%FCS in DMEM for 2–4 hr before cell plating. Then the medium from different inlets flow following a programmed sequence through the channels around the microfluidic wells and diffuse through a perfusion barrier protecting the cells, minimizing the undesirable effect of shear stress. The flow rate obtained of 10 μl/h across the small volume of the well (less than 1 μl) allows a replacement of the medium in contact with the cell within minutes.

### Quantification and Statistical Analysis

#### Automated analysis of smFISH data

The smFISH data displayed in Figure 1 were analyzed using the Matlab-based software FISHquant (Mueller et al., 2013). Mature RNA were identified as 3D gaussian spots with peak intensity above an arbitrary threshold, which was kept constant for all the stacks belonging to the same RNA specie. Nascent RNAs at active TS were quantified by identifying the sites of nascent transcription as bright nuclear foci, setting the threshold so that no more than four actively transcribed loci could be found within each nucleus. For each of the transcription sites the amount of RNA was calculated by comparing the integrated intensity of the site with the average integrated intensity of the spots identified as mature RNAs. Active TS displaying less than two transcripts were filtered out from subsequent analysis as they are practically indistinguishable from released mature. The smFISH stacks are displayed as maximal projections. Distributions of mature RNAs in Figure S1C were fit via a negative binomial model that – under the assumption of the random telegraph model (Raj et al., 2006) - provide the probability of observing a certain number of RNAs per cell *P*(*x*) as function of the relative burst frequency *f_rel_* and burst size *b* as:

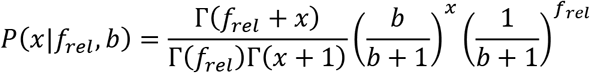

Details on the statistical analysis are reported in the relevant figure legends.

#### Automated analysis of live-cell imaging data

To study TS activity dynamics, time lapse of the z-stacks are maximal-projected and filtered. The nuclei are segmented using the MCP-GFP signal and using a low-pass filter the TS are detected as bright spots (red arrows in **Supplementary Figure S3_1 (A)**). The information is combined to track the TS within each cell and to obtain the TS signal dynamics for hundreds of cells. To quantify NF-κB nuclear localization dynamics in living cells, we use the same nuclear mask as for the MCP-GFP signals and quantify the nuclear concentration of NF-κB normalized by the cytosolic fluorescence intensity (Ashall et al., 2009; Nelson et al., 2004; Zambrano et al., 2016). Responding cells are those reaching a nuclear to cytosolic NF-κB ratio greater than one. Routines are available upon request and run on Matlab R2015.

#### Stochastic fitting and mathematical modelling

Detailed information is provided in the **Supplementary Methods**.

##### Stochastic fitting

Briefly, we adapted our algorithms (Molina et al., 2013; Suter et al., 2011) to compute the likelihood of the time traces obtained, considering the TS signal and the standard deviation of the background, compared to those generated by a simple stochastic gene expression telegraph-like model. In such model, the gene can switch between and active and inactive state and transcription occurs in bursts only during the active periods. MCMC sampling from the posterior distribution was performed to estimate to infer gene activity, the signal-to-transcripts scaling, the rates of accumulation and release of new transcripts *k*^+^ and *k*^−^ and the rates of the gene activation process k_on_ and k_off_. Calibration was performed by imposing an average value of the number of nascent transcripts at t=20 min equal to the value observed by smFISH. *Deterministic mathematical model of NF-kB mediated gene activation* was performed using a simple model of NF-kB dynamics using the core negative feedback of the system. ODE simulations were performed using matlab. *Stochastic simulations:* we developed a C++ software, *hysim*, able to run stochastic simulations using the Gillespie algorithm. We used such software to generate trajectories of the stochastic version of the biochemical networks analyzed using the deterministic approach. We used a similar approach to obtain nascent transcription simulation taking as an input the NF-κB nuclear localization dynamics in single cells.

## Supplementary Information

**Table S1, related to Figure 1.** List of smiFISH probes used in the paper.

### Supplementary Methods

**Movie S1, related to Figure 3.** Exemplary time-lapse acquisition for cells treated with 10ng/ml TNF-α. Shown are maximal projections.

**Movie S2, related to Figure 3** Exemplary time-lapse acquisition for untreated cells. Shown are maximal projections.

**Movie S3, related to Figure 3.** Exemplary single-cell analysis of the MS2 signal intensity in a single cell upon treatment with 10ng/ml. The displayed cell shows a prompt response in minutes upon stimulation.

**Movie S4, related to Figure 3.** Exemplary single-cell analysis of the MS2 signal intensity in a single cell upon treatment with 10ng/ml. The displayed cell shows a late response.

**Movie S5, related to Figure 3.** Exemplary single-cell analysis of the MS2 signal intensity in a single cell upon treatment with 10ng/ml. The displayed cell shows no response.

**Movie S6, related to Figure 4.** Exemplary time-lapse acquisition for cells treated with two pulses of 10ng/ml TNF-*α* as described in figure 4. Shown are maximal projections.

**Movie S7, related to Figure 5.** Exemplary simultaneous acquisition of NF-kB translocation (left) and MS2 transcription dynamics (center, maximum projection shown). The overlay of the two channels is also shown.

**Movie S8-S10, related to Figure 5.** Exemplary single-cell analysis of the MS2 signal intensity (left, displayed in green in the plot) and NF-kB translocation (center, displayed in red in the plot) in a single cell upon treatment with 10ng/ml.

**Movie S11, related to Figure 6.** Exemplary time-lapse acquisition for cells treated with TNF-*α*+CHX. Shown are maximal projections.

## Supplementary methods of “First responders shape a prompt and sharp NF-**κ**B-mediated transcriptional response to TNF-*α*” by Zambrano et al. (2020)

### A. Stochastic model for parameter inference

To estimate transcriptional kinetic parameters and gene activity patterns of single cells, we developed a stochastic model of transcription combined with a Bayesian inference approach similar as in [1]. Briefly, we described MS2 locus as a stochastic system that is characterized by two random variables, the gene state *g* that indicates whether the gene is transcriptionally active (*g* = 1, *G_on_* in the main text) or inactive *g* = 0 (*G_off_* in the main text) and the number of nascent transcripts *n* on the TS. Thus, transcription at the MS2 locus can be described as a stochastic process emerging from the following set of biochemical reactions:

1. Gene activation and deactivation: 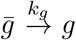, where *k*_1_ is the activation (*k_on_* in the main text) rate when the gene is inactive (*g* = 0) and *k*_0_ (*k_off_* in the main text) is the inactivation when the gene is active (*g* = 1).
2. Linear increase in the number of transcripts on the TS: 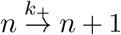.
3. Linear decrease in the number of transcripts on the TS: 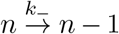, where *k_e_* is an effective elongation rate.

Note this model is a concise simplification of transcription where many complex processes are described by single effective reactions. Indeed, the rates *k_on_* and *k*_+_ summarize chormatin remodeling, transcription factor binding, preinitation complex formation and RNA polymerase initiation. In turn, *k*_−_ represents transcript elongation, splicing and termination. Importantly, we chose linear increase and decrease of the number of transcripts based on the results shown in Ref. [2]. In short, since RNA polymerases travel in convoys, the transcription site (TS) displays peaks of intensity that increase and decrease linearly, and whose slopes depend on the elongation rate and on the interspace between polymerases. Finally, we assumed that the MCP binding/unbinding process to the MS2 loops is fast compared to the elongation of transcripts and therefore is not explicitly modeled. In spite of all these simplifications, the model is able to accurately fit the data and provide information about the dynamics of gene activation/inactivation.

The chemical master equation describing the stochastic dynamics of the system can be written as,

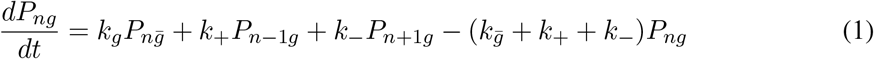

which, truncating the system at a sufficiently large number of transcripts *n*_max_ = 100, can be expressed in matricial form,

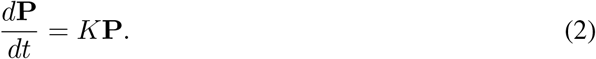

This truncated master equation is then a linear system of ordinary differential equations and therefore can be easily solved numerically by calculating the exponential of the rate matrix *K*. Thus the propagator of the stochastic system is obtained as:

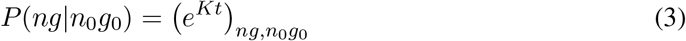

which describes the transition probability from a initial state (*n*0, *g*0) to a final state (*n*, *g*) in a given time *t*.

#### A.1. Noise model

The next key ingredient for our inference approach is to introduce a noise model that relates the state of the system at given time with the measured fluorescence signal. A simple but reasonable and convenient choice is to assume that the expected amount of signal *s* is proportional to the number of nascent transcripts *n* plus a background signal level *b*. Furthermore, we assumed that the fluctuations around the expected mean are Gaussian distributed with a standard deviation *σ*. Under these assumptions the noise model can be expressed as,

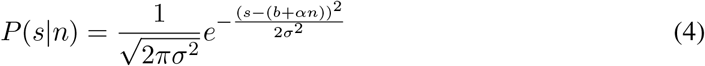

where *α* is the scaling factor that relates the number of nascent transcripts with the expected observed fluorescence signal. Equation 4 can be considered as the emission probability, i.e. the probability that the system emits a signal *s* given that is in the state *n*. Note again that this simple model does not take into account the MCP dynamics which can be an additional source of noise.

#### A.2. Inference

The propagator of the system and the noise model introduced above allow us to calculate the probability of observing an experimental time series consisting of *N* measurements of MS2 signal *S* = {*s*_1_, *s*_2_, …, *s_N_*} given the model parameters Θ = {*α*, *σ*, *k_g_*, *k*_+_, *k*_−_}. Indeed, this probability can be expressed as the product of the probability of the signal *S* given that the system went through a particular state trajectory times the probability of that trajectory and then summing over all possible trajectories, i.e:

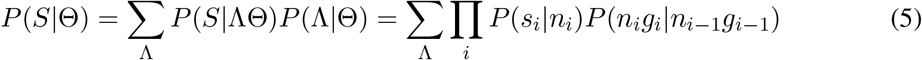

Importantly, the noise model and the propagator can be considered as emission and transition probabilities of a Hidden Markov Model and therefore we can use linear programming to efficiently sum over all possible state trajectories [1]. Then, assuming that cells are independent of each other, the probability of observing the signal of *C* cells *D* = *S*_1_, *S*_2_, …, *S_C_* can be written as:

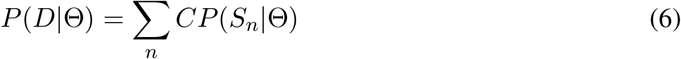

Then, applying Bayes’ theorem we can obtain a posterior distribution over the parameters Θ given the data *D*:

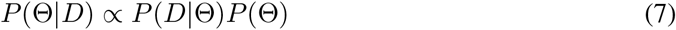

where we used a scale invariant prior for the parameters, i.e *P*(*θ*) = 1/*θ*. Finally as the posterior probability over the parameters cannot be calculated analytically we used Markov Chain Monte Carlo (MCMC) to sample it and to obtain average and standard deviations for each parameter.

Once the model parameters are estimated we can again apply Bayes’ theorem to obtain a probability distribution over the hidden state trajectories given a time series of MS2 signal *S*:

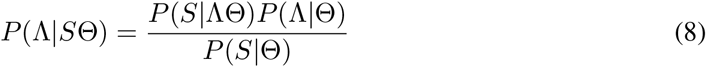

Notice that the number of possible trajectories grows exponential with the number of data points. However, we can use HMM tools to calculate efficiently the trajectory Λ* that maximize the distribution [1].

In conclusion using stochastic modeling of biochemical reaction in combination with a Bayesian inference approach we were able to estimate effective transcriptional parameters and temporal profiles of gene activity.

#### A.3. Statistical independence of the response to consecutive pulses of TNF-*α*

Using the approach above it is possible to estimate the periods of gene activation/inactivation within certain time interval. In particular, in our experiments in which 1 hour 10 ng/ml TNF-*α* pulses are followed by 2 hours washouts we can estimate the probability of having a gene activation in certain time intervals after each TNF-*α* pulse.

We can call *p*_1_ to the fraction of cells displaying a burst at most 2 hours after the beginning of the first TNF-*α* pulse, and *p*_2_ to the fraction of cells displaying a burst at most 2 hours after the beginning of the second pulse. Both can be readily estimated from our data. If both events are independent, the fraction of cells displaying no bursts should be (1 – *p*_1_) · (1 – *p*_2_), those with a burst only after the first TNF-*α* pulse should be *p*_1_ · (1 – *p*_2_), while the fraction of cells displaying a burst only after the second TNF-*α* pulse should be (1 – *p*_1_) · *p*_2_. Finally, the fraction of cells with a burst after each TNF-*α* pulse is *p*_1_ · *p*_2_. We calculated such theoretical distribution for three independent experiments and found that it clearly departs from the distribution observed in the experiments.

### B. Deterministic and stochastic modelling NF-**κ**B - mediated transcription

#### B.1. Model of the NF-**κ**B system

For our qualitative exploration of NF-**κ**B mediated transcription we used a simple model of the NF-**κ**B system able to recapitulate the essential features of NF-**κ**B nuclear localization dynamics upon TNF-*α* [3]. In such model for simplicity it is considered that we can either have free NF-*κ*B (hence nuclear and transcriptionally active) or forming a complex with the inhibitor I*κB*, NF-*κ*B:I*κ*B (the cytosolic and transcriptionally inactive form). We represent their copy number as *NFκB* and *NFκB* : *I*κ*B*. The total amount remains unchanged, so *NF*κ*B* + *NF*κ*B* : *I*κ*B* = *NFκB_tot_*. An external signal (for us, TNF-*α*) can lead to the activation of a kinase complex that leads to the degradation of the inhibitor and hence sets free (and active) NF-*κ*B. For simplicity we assume that an external stimulus produces instantaneously a constant number of active kinase *IKK*_0_>0 in presence of external stimulus, and *IKK*_0_=0 for unstimulated cells. When NF-*κ*B is free, it can activate the genes encoding for the inhibitor, that would go from inactive *G_I,i,off_* to active *G_I,i,on_*, producing the transcript I*κ*B_*RNA*_ that is then translated (*i* = 1, 2 stands for each of the alleles). The copy numbers of the transcript and the inhibitor protein are written below as *IκB_RNA_* and *I*κ*B* respectively.

The deterministic model for this system [3] can be written as:

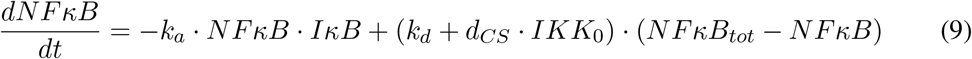

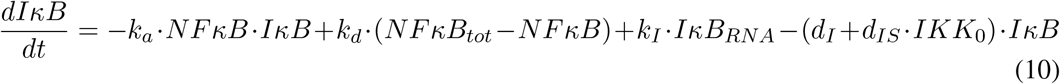

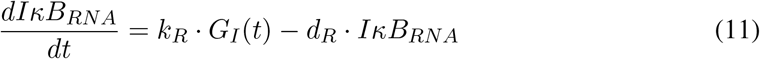

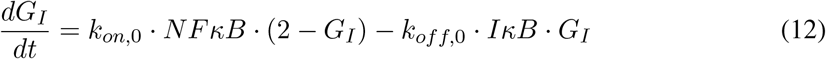

where *G_I_*(*t*) = *G*_1;*on*_(*t*) + *G*_2,*on*_(*t*). The values of the parameters are provided in table B.1

**Table B.1:** Parameters for the NF-*κ*B system

| Name | Value | Units |
| --- | --- | --- |
| $IKK_0$ | $10^5$ | mols |
| $NF\kappa B_{tot}$ | $310^4$ | mols |
| $k_a$ | $1.9 \cdot 10^{-6}$ | $\text{mols}^{-1} \cdot \text{s}^{-1}$ |
| $k_d$ | $8.4 \cdot 10^{-5}$ | $\text{s}^{-1}$ |
| $d_{CS}$ | $2.5 \cdot 10^{-8}$ | $\text{mols}^{-1} \cdot \text{s}^{-1}$ |
| $k_I$ | $2.5 \cdot 10^{-1}$ | $\text{s}^{-1}$ |
| $d_I$ | $6.7 \cdot 10^{-5}$ | $\text{s}^{-1}$ |
| $d_{IS}$ | $5 \cdot 10^{-9}$ | $\text{mols}^{-1} \cdot \text{s}^{-1}$ |
| $k_R$ | $2 \cdot 10^{-1}$ | $\text{mols} \cdot \text{s}^{-1}$ |
| $d_R$ | $7.5 \cdot 10^{-4}$ | $\text{s}^{-1}$ |
| $k_{on,0}$ | $6.9 \cdot 10^{-8}$ | $\text{s}^{-1}$ |
| $k_{off,0}$ | $1.4 \cdot 10^{-8}$ | $\text{s}^{-1}$ |

#### B.2. Deterministic, stochastic and hybrid simulations

To simulate the intrinsic variability of the system, an alternative is to take the biochemical reactions that give rise to the mass action kinetic equations described above (details provided in [3]) and to perform stochastic simulations using e.g. the Gillespie algorithm. However a less time consuming approach is to perform what we can call hybrid simulations, in which the evolution in time of the variables with high copy numbers are modelled using ordinary differential equations while those with low copy numbers are modeled using an approximation of the next-reaction method (full description is provided in the appendix). This is indeed the approach that was followed in other works to model variability of the NF-*κ*B nuclear localization dynamics [4–6], and applied to our model this would imply to substitute Eq. 12 by the following stochastic process:

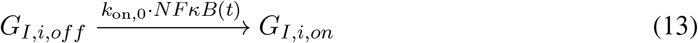

and of inactivation

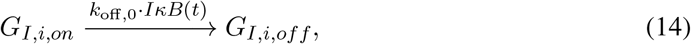

where *i* = 1, 2.

**Figure B.1:**
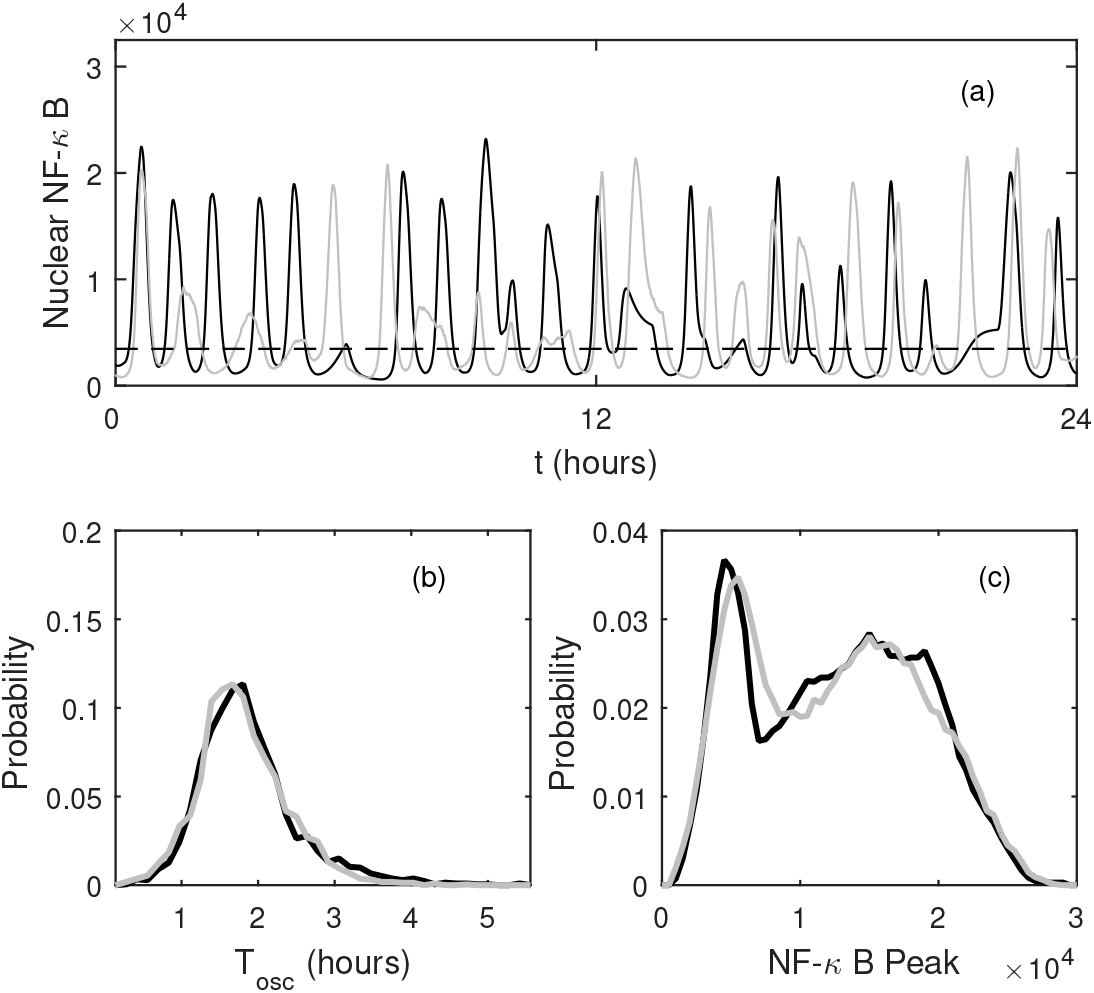
Results in each panel were obtained using *hysim* to perform stochastic simulations (gray line), hybrid simulations where only the gene activation/inactivation is modelled stochastically (black line) and deterministic simulations (—) (a) Example of sustained oscillations obtained for the stochastic and hybrid simulations, but not for the deterministic. For hybrid and stochastic simulations we obtained the distribution of the oscillatory periods *T_osc_* that peak at about 1.5 hours and (c) of the peak values of each oscillation. The distributions of the hybrid simulations that we and others use for simulating NF-*κ*B signalling and fully stochastic simulations give fairly similar results.

To our knowledge a comparison between fully stochastic simulations and this “hybrid” has never been performed. We developed a software called *hysim* that one can use to flexibly decide which variables shall be modeled as deterministic and which as stochastic processes. By using it, we can say that both do provide a fairly similar sustained oscillations (as opposed to the purely deterministic model, see Fig. B.1(a)). More importantly, the distributions of the peak values and the peak periods Fig.B.1 (b) and (c), which justifies the use of the hybrid approach for stochastic simulations.

#### B.3. Models of NF-*κ*B-mediated transcription of target genes

We used the mathematical model of the NF-*κ*B system described above as an input for the activation dynamics of a target gene *G* that can switch between *G_on_* and *G_off_* states following three schemes. In all of them, following considerations on the non-cooperativity of NF-*κ*B mediated gene activation [7], we consider that the gene activation probability depends linearly on the nuclear concentration of NF-*κ*B:

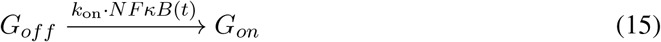

The models considered differ in their inactivation rates.

For the model i whe have that the inactivation is spontaneous, of the form:

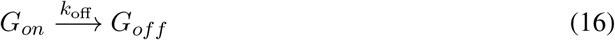

Instead, for model ii we consider that the inactivation is mediated by molecular stripping:

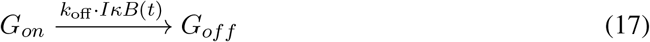

Finally, in model iii we consider a refractory state, meaning that the inactivation is spontaneous but leads to a refractory state *G_ref_* so

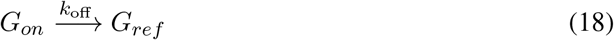

and

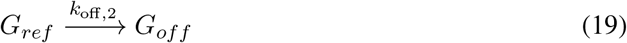

**Table B.2:** Reference values of the parameters for the models of NF-*κ*B mediated transcription

| model | Name | Value | Units |
| --- | --- | --- | --- |
| i,ii,iii | $k_{on,0}$ | $6.9 \cdot 10^{-8}$ | $\text{mols}^{-1} \cdot \text{s}^{-1}$ |
| i,iii | $k_{off,0}$ | $3.7 \cdot 10^{-4}$ | $\text{s}^{-1}$ |
| ii | $k_{off,0}$ | $1.4 \cdot 10^{-8}$ | $\text{mols}^{-1} \cdot \text{s}^{-1}$ |
| iii | $k_{off,2,0}$ | $3.7 \cdot 10^{-4}$ | $\text{s}^{-1}$ |

The reference values around which we perform our numerical exploration (for up to two orders of magnitudes above and below) for each model are specified in table B.2. They are based on those used in our model of the NF-*κ*B system, which were themselves derived from the literature (details in Ref.[3]). Since models i and iii do not contemplate stripping, their effective value of reference inactivation rate (*k*_*off*,0_) is equal to the one of model ii (with stripping) multiplied by the average I*κ*B levels 3 hours post-stimulation.

To explore the promptness and the sharpness of the gene response at population level we used fully deterministic simulations of the above equations, which imply to add to the set of equations 9 to 12 the following ones for the gene activity:

- For Model i:

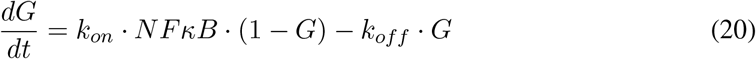
- For Model ii:

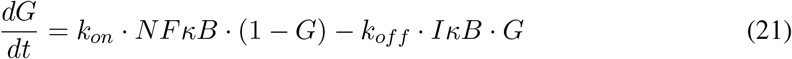
- For Model iii:

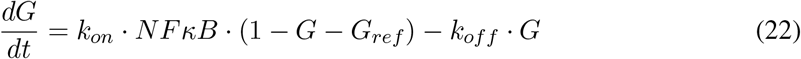

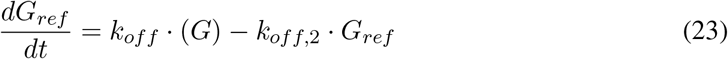

Instead, for stochastic simulations we used as an input for the stochastic processes above the hybrid simulations of the NF-*κ*B system. In all of the models, we allowed the number of nascent transcripts *n* to grow and decrease incrementally following the equations:

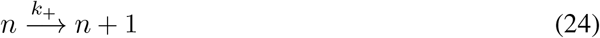

and decreasing incrementally

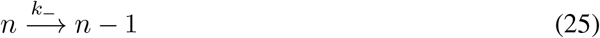

Where we used values of *k*_+_ and *k*_−_ producing bursts of fast increase and decrease of the signal as compared to NF-*κ*B translocation dynamics.

Finally, to ensure that the observed result did not depend on the particular shape of the NF-*κ*B nuclear localization dynamics, performed simulations of the model iii using as an input the experimentally obtained data of the NF-*κ*B nuclear localization dynamics in single cells. In short, we used the data to compute the integrals necessary for application of the Gillespie algorithm, such as Eq. 32 (see Appendix) and produce simulations of nascent transcription for single cells and averaged across the population.

### C. Search time calculation for NF-*κ*B targets

In a recent paper [8], we applied single molecule tracking (SMT) to quantify the NF-*κ*B binding kinetics at specific and at non-specific binding sites in HeLa cells. Upon stimulation with TNF-*α*, NF-*κ*B displayed a bound fraction equal to approximately 20 %. Mutant analysis allowed to identify that NF-*κ*B bound molecules partitioned into a transient non-specifically bound population ((*f_ns_* = 96%,*τ_ns_* = 0.5 s) and a more stable population representing specific binding *f_s_* = 4%,*τ_s_* = 0.4 s) As described in [9] we can use these quantities to estimate the time that it takes for a single NF-*κ*B molecule to reach one of its specific targets. The average residence time of NF-*κ*B on chromatin can be calculated as:

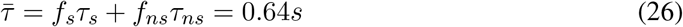

From this, we can then estimate the average free-diffusion time between two binding events as:

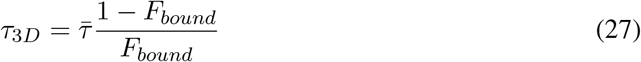

The search time to find a specific site can then be obtained by knowing the number binding events that a molecule needs to undergo on average before encountering a specific binding site: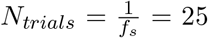. Each trial round will take a time equal to *τ*_3*D*_ + *τ_ns_*, except for the last one which will last *τ*_3*D*_, after which a specific site is found. We can therefore calculate the search time as:

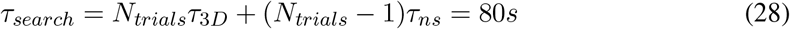

This is the average time that it takes for one single NF-kB molecule to find one of its target sites. By dividing *τ_search_* for published estimates [10] of NF-*κ*B molecules (approx. 30000) and by multiplying it for the number of NF-*κ*B target sites (estimated by the number of high-confidence peaks obtained by ChIP-seq on NF-*κ*B [11] (approx. 50000) we can provide a rough estimate of approximately 2 minutes for the time that it will take for a specific NF-*κ*B target gene to be found by any of the available NF-*κ*B molecules.

### D. Stochastic simulations with *hysim*

The hybrid simulation approach proposed was described in detail [12], and is a simplified version of the approach described in [13] to study the evolution in time of a system of biochemical species with a wide variety of copy numbers and reaction speeds. The formal description of the approach would be as follows: consider that the state of this system in time *t* is determined by the vector state *X* = (*X*_1_(*t*),*X*_2_(*t*), …, *X_N_*(*t*)), where *X_j_*(*t*) is the number of copies of the biochemical species *j* at time *t*. Such species can interact through *M* biochemical reactions with rates *a_j_*(*X*(*t*)), so the probability of the *j*-th reaction taking place in *dt* is *a_j_* (*X* (*t*))*dt*, while *v_ji_* denotes the change in species *i* due to the *j*-th biochemical reaction.

In [13], in order to speedup stochastic simulations, it is proposed to approximate as a Langevin equation the evolution of the variables that satisfy the two following conditions

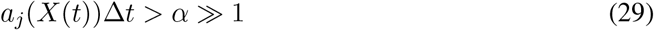

and

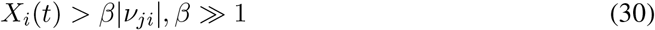

where the first equation imposes that many reaction events will take place in Δ*t*, while the second ensures that the number of molecules is much bigger than the change in the number of molecules caused by the reaction. The bigger *α* and *β* are, the better the approximation to the Langevin equation is. The remaining species of the system should be modelled as a Markov process, using for example the Gillespie algorithm [14].

In the case of stochastic simulations of genetic circuits, it is clear that certain biochemical species will not satisfy eqs. 29 and 30: for example, the maximum number of active genes encoding for a given protein is (typically) two, so eq. 30 will not be satisfied. The remaining biochemical species have copy numbers that go from *O*(100) for the transcripts to *O*(10^5^) for the proteins [15] so, if eqs. 29 and 30 are fulfilled, they could be modelled using a Langevin equation. In this case, since the relative fluctuations of these variables shall be small, an even simpler approximation would be to model the dynamics of variables satisfying eqs. 29 and 30 using ordinary differential equations instead of the proposed Langevin equation [13]. This idea was used in different works dealing with NF-*κ*B dynamics [4–6] and it is the same idea that we applied for our simulations of a simple model of the NF-*κ*B system.

In order to apply this approach to our model and (in principle) to any other models of signaling pathways through mass action kinetics equations we created a sofware in c++, *hysim*, that performs hybrid and fully stochastic simulations for an arbitrary biochemical system of reactions by selecting which variables of the system should be modeled deterministically and which stochastically. The *hysim* software can be downloaded at https://github.com/MolinaLab-IGBMC/hysim, where we also provide a userguide.

To describe *hysim*, we can generalize the hybrid integration scheme by defining the *deterministic variables* vector as *D*(*t*) = (*X*_1_(t), *X*_2_(*t*), …, *X_D_*(*t*)) and the *stochastic variables* vector *S* = (*X*_*D*+1_(*t*), *X*_*D*+2_(*t*), …, *X_N_*(*t*)). A possible criterion for this would be to choose for *D* the variables that satisfy 29 and 30, and leave the remaining for *S*. Without loss of generality, we can say that reactions going from *n* = 1 to *n* = *N_S_* ≤ *M* are those that imply a change in the number of copies of some of the stochastic variables, i.e. that for 1 ≤ *n* ≤ *N_S_*, *v_ni_* ≠ 0 for some *D* + 1 ≤ *i* ≤ *N*.

In this situation, our hybrid modelling scheme implies that the system evolves in time as prescribed by

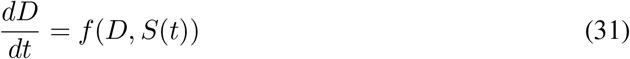

where the variables of the vector *S*(*t*) would evolve stochastically as a Markov process, according to the next-step Gillespie algorithm [14].

In other words, given the state of the system at time *t*, we generate two random numbers *r*_1_ and *r*_2_ in the [0,1] interval and find the time *τ* at which the next reaction takes place [14], for which

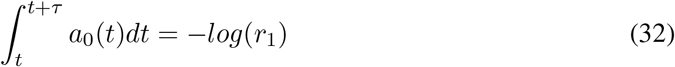

where *a*_0_ is the cumulative probability of a reaction including a stochastic variable taking place, i.e.

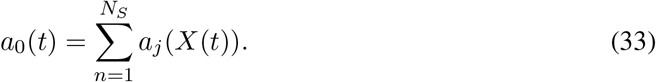

Once *τ* is found, the reaction that takes place is selected by choosing the *k* such that

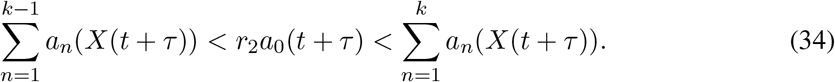

Once the change is found the vectors *X*(*t*) and *S*(*t*) are updated as prescribed by *v_ki_*, and the process can be repeated for as long as required. This procedure speeds up simulation consistently as compared to fully stochastic simulations while giving similar results (see the examples above).

---

**Figure S1, related to Figure 1.**
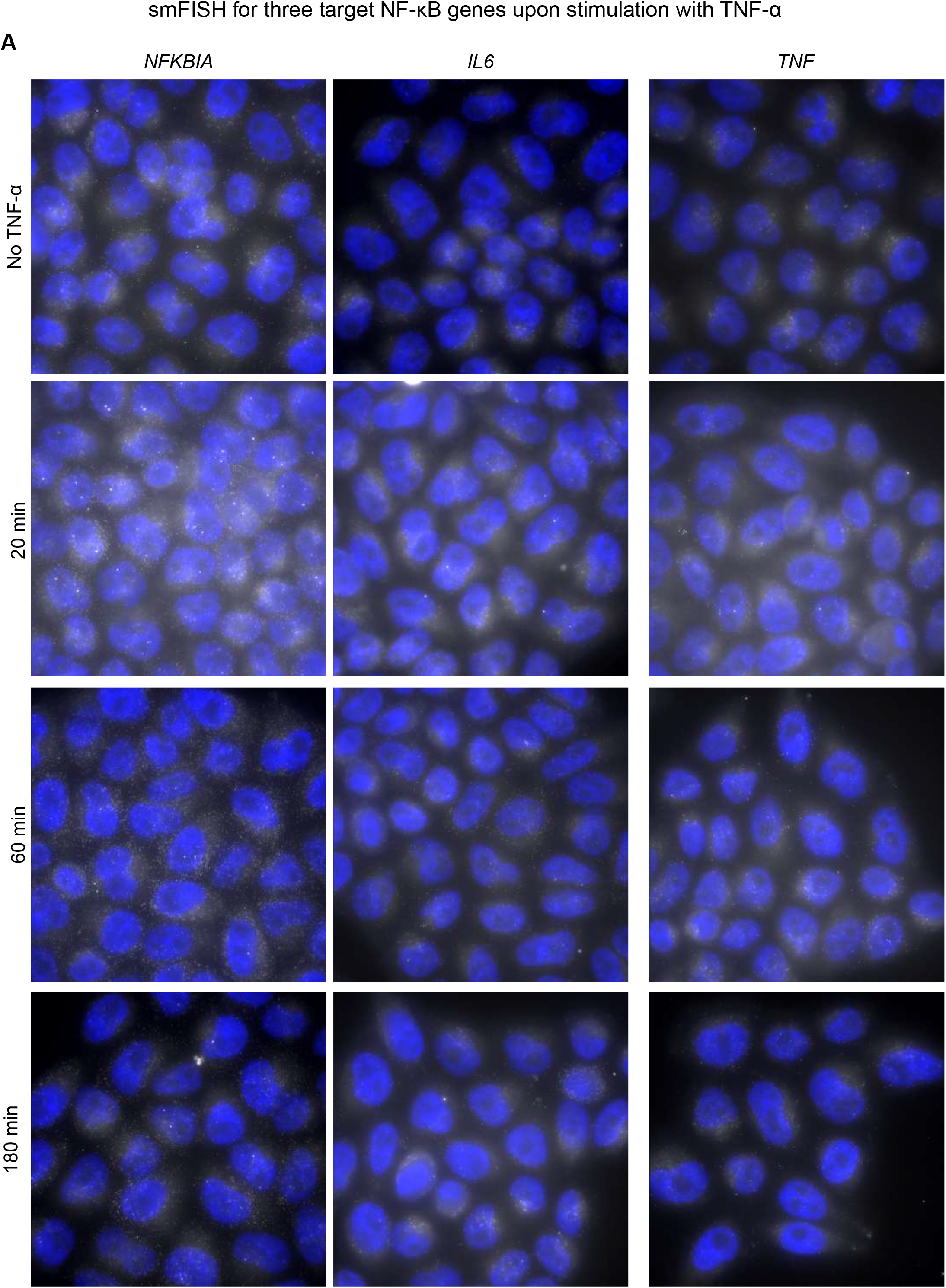
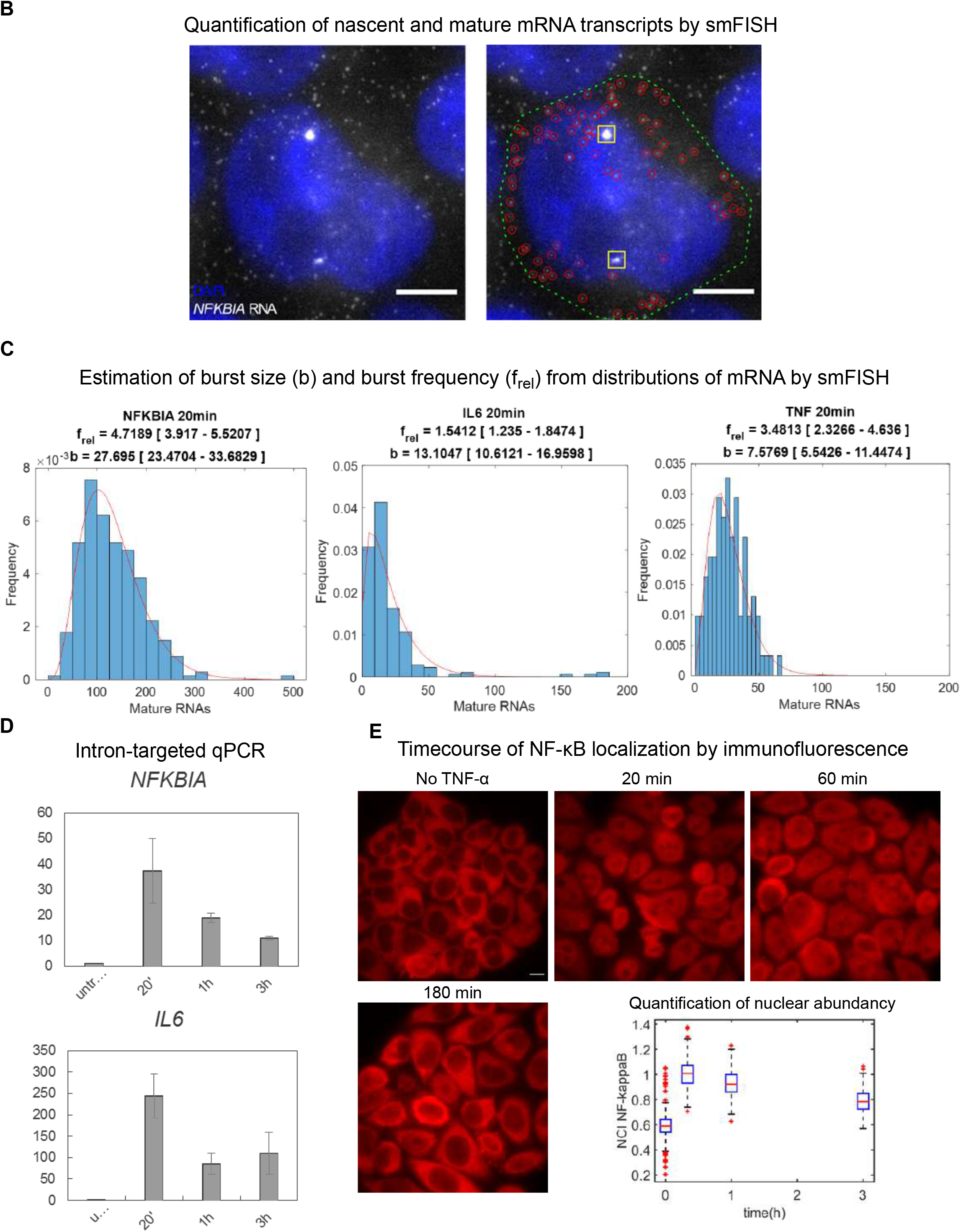
**A.** Exemplary fields of smFISH acquisitions for the three different targets at different time-points. Maximum projection displayed. **B.** A cell displaying an active transcription site (TS) and single RNAs, detected using the smFISH quantification software. **C.** Fitting of the distribution of mature RNAs obtained by smFISH with a negative binomial allow estimating the relative frequency of transcriptional bursts *frel* and the burst size *b*. **D.** Nascent RNA estimated by qPCR for two NF-kB targets upon TNF-α, also reproduces the average dynamics described in our single-cell assays. **E.** Immunofluorescence against NF-κB for our cells stimulated with 10 ng/ml TNF-α and quantification using our routines, showing a peak at 20 minutes.

**Figure S2, related to Figure 2.**
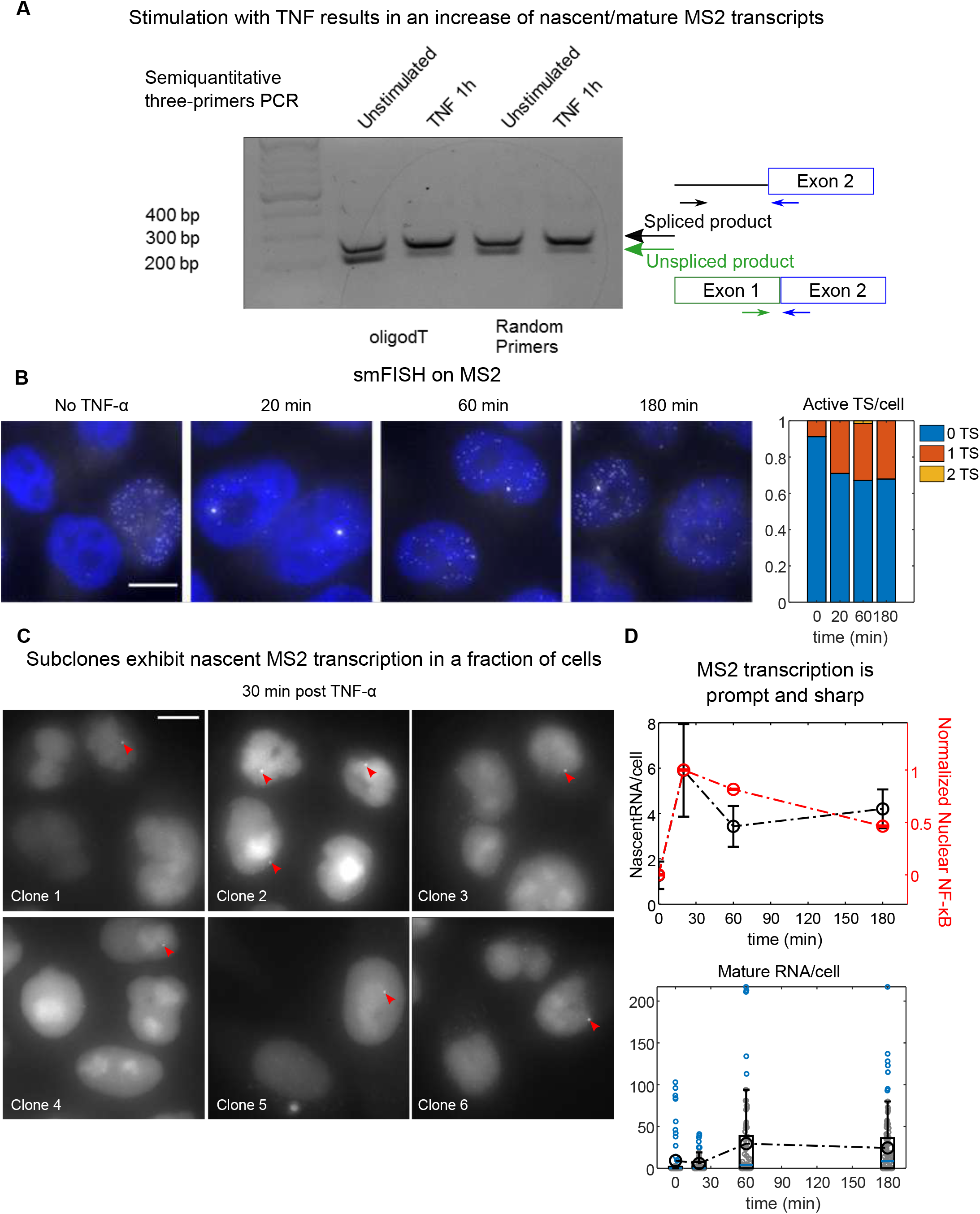
**A.** PCR products of the unspliced (actively transcribed) and spliced form of the reporter’s RNA product, showing a shift towards unspliced form in TNF-α stimulated cells after 1 hour, which indicates active transcription. **B.** Exemplary smFISH acquisitions using probes targeting *MS2* RNA at different times following TNF-α and fraction of cells with either 0,1 or 2 active transcription sites. **C.** Representative mages of six out of 10 clones generated from our cells for which at least 1 clear active TS per image is observed (red arrow). **D.** Average number of nascent MS2 transcripts per cell measured by smFISH (black, error bars SEM, n_cells_ =80, 79, 76, 106 for 0’, 20’, 1 hour, 3 hours time -points respectively) and normalized to nuclear-to-cytosolic NF-κB fluorescence intensity assessed by immunofluorescence (red, errorbars, SEM n_cells_ =326, 225, 212, 211 for 0’, 20’, 1 hour, 3 hours time-points respectively). Nascent MS2 RNA peaks at 20’ and decays faster than the NF-κB nuclear abundancy.

**Figure S3_1, related to Figure 3.**
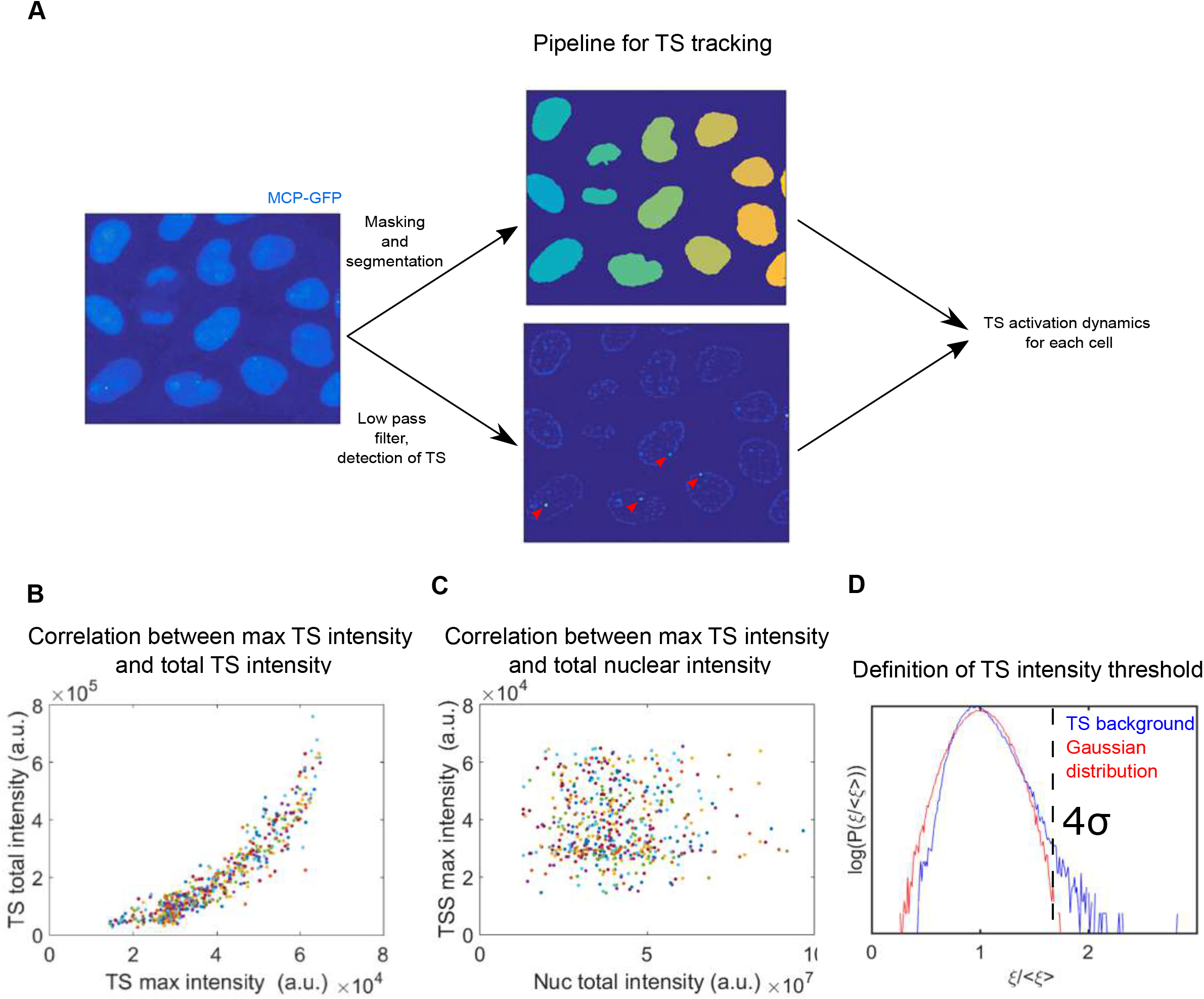
**A.** Workflow of the live cell imaging approach to study TS activity dynamics. Time lapse of the z-stacks are maximal-projected and filtered. The nuclei are segmented using the MCP-GFP signal and using a low-pass filter the TS are detected as bright spots (red arrow). The information is combined to track the TS within each cell and to obtain the TS signal dynamics for hundreds of cells. **B.**Correlation of the TS max intensity and the total TS intensity. Each dot corresponds to a single cell’s time series (Pearson’s correlation coefficient r ^2^=0.84). **C.** Absence of correlation between the TS max intensity and the nuclear intensity (r^2^=5·10^−5^). Each dot corresponds to a single cell’s time series.**D.**Probability distribution function of the average -normalized backgrou nd signal around the detected TS (blue line) compared to a normal distribution with the same mean and standard deviation (red line). The graph show that the distribution is long-tailed so *p(ξ>μ*+*4s*)=2·10^−3^. When the maximum value of the detected TS is beyond the threshold, the gene is considered “active”; with this threshold the probability of observing a false positive in 60 frames is below 5%.

**Figure S3_2, related to Figure 3.**
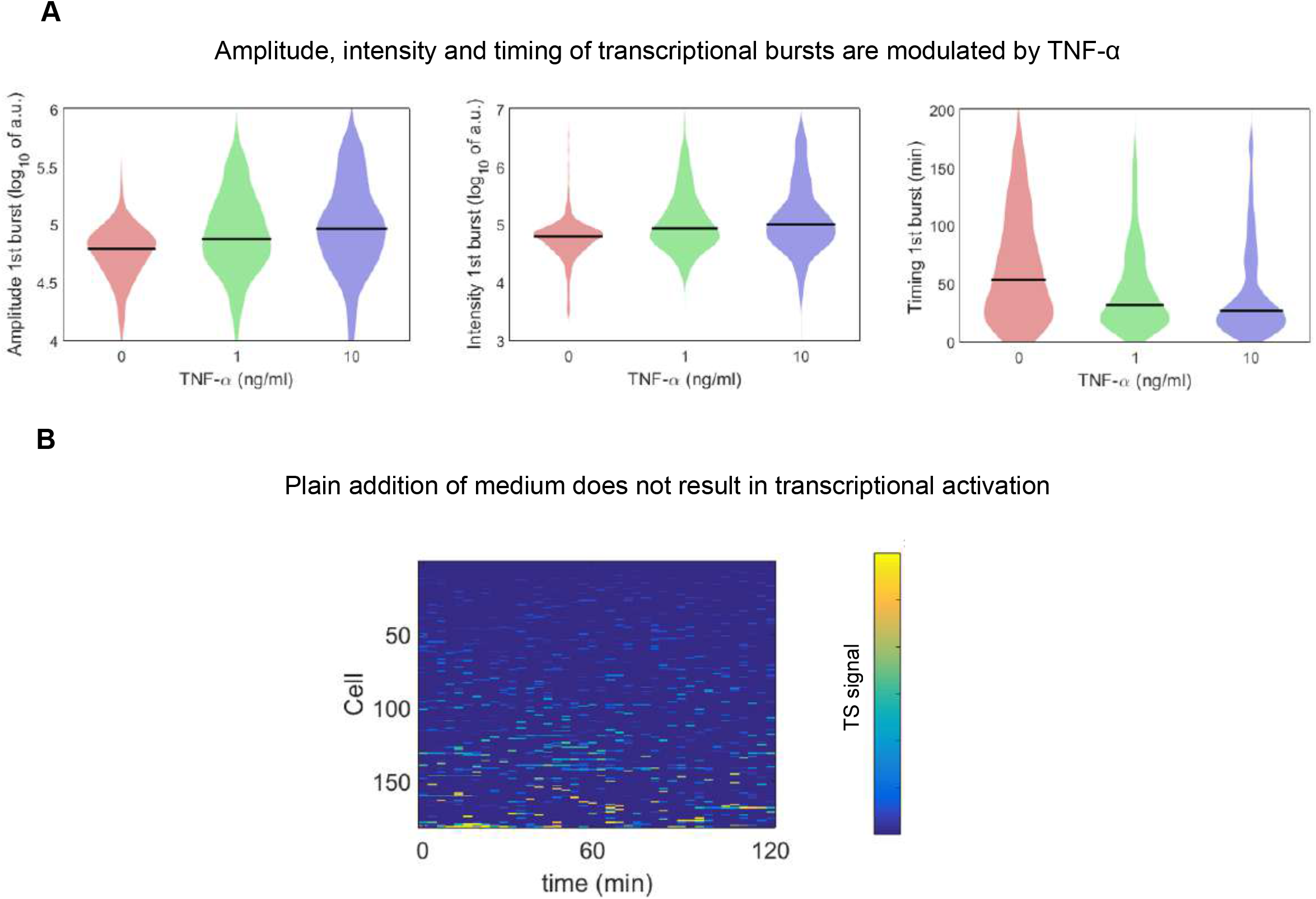
**A.** Features of the detected transcriptional bursts from three representative experiments for each TNF-α dose considered. **B.** TS signal for cells to which plain medium was added, confirming that the medium addition itself does not lead to significant TS activation.

**Figure S3_3, related to Figure 3.**
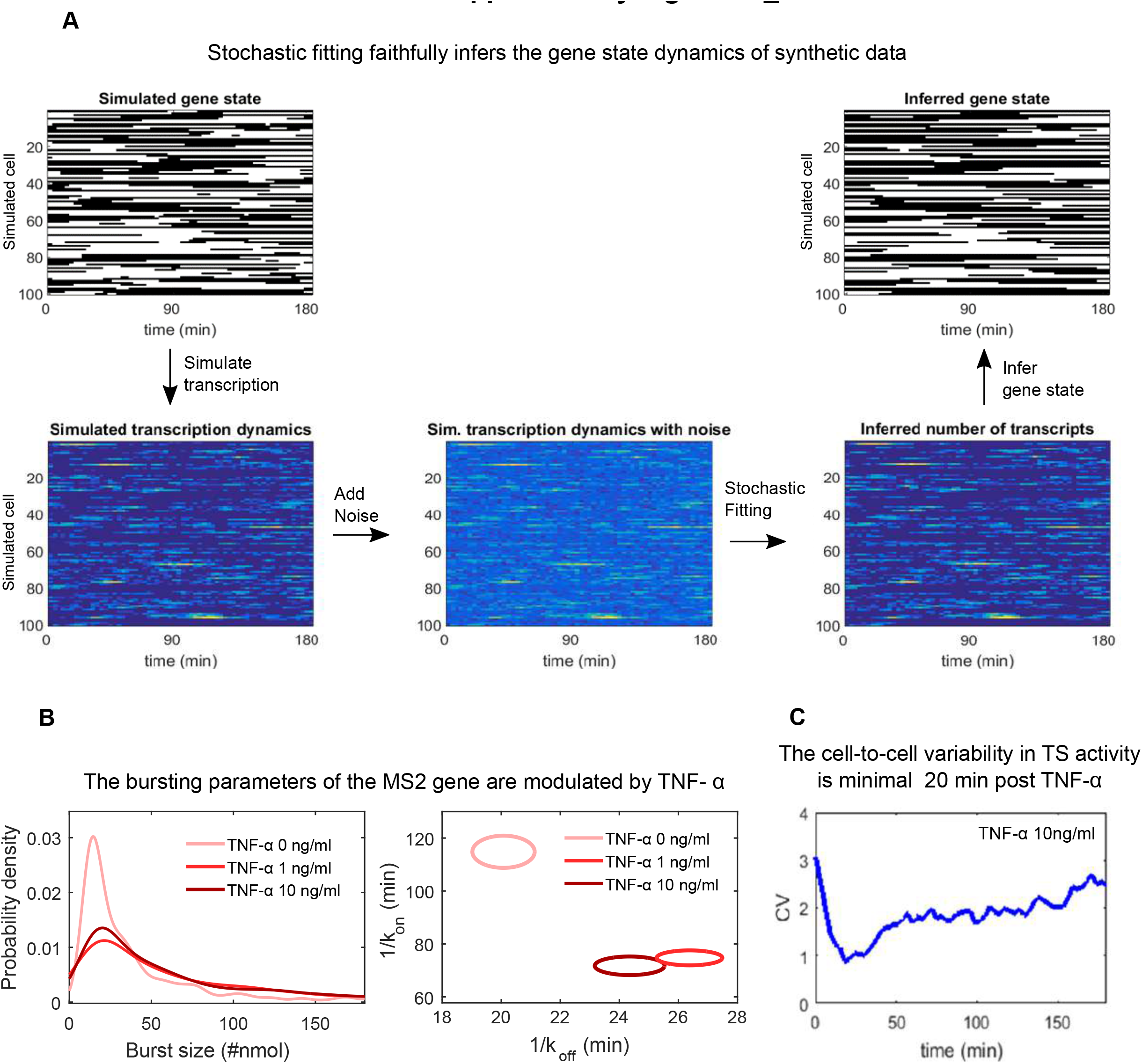
**A.** Example of single-cell traces generated from a stochastic model of activation-inactivation, on top of which our model of transcription initiation and elongation is superimposed given rise to bursts. Noise was added to the signal and then we applied our inference model to deconvolve the signal, obtaining inferred nascent transcript and gene activity signals that faithfully reproduce the original ones. **B.** Distributions of burst-sizes and parameters obtained from our fittings of the experiments from the three conditions considered. **C.** Coefficient of variation (ratio of standard deviation and mean) of the “prompt responders” as a function of time, showing a decrease at approximately 20 mins post-stimulation and a return to basal variability levels.

**Figure S4, related to Figure 4.**
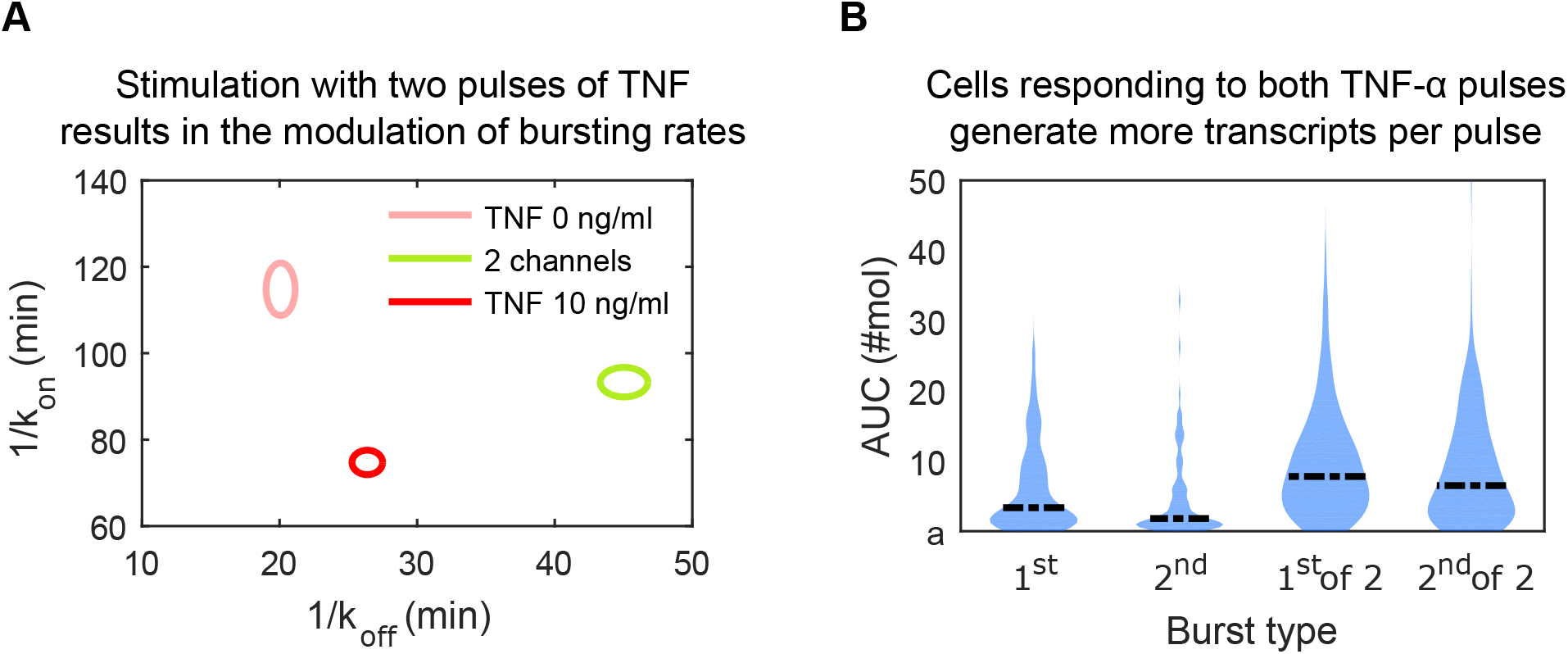
**A.** Parameters obtained by the stochastic fitting of the experiments obtained upon stimulation with two TNF-α pulses. **B**. Transcriptional activity, inferred as area under the curve (AUC) following the first and the second TNF-α pulse for cells responding to only one of the two pulses, or to both of them.

**Figure S5, related to Figure 5.**
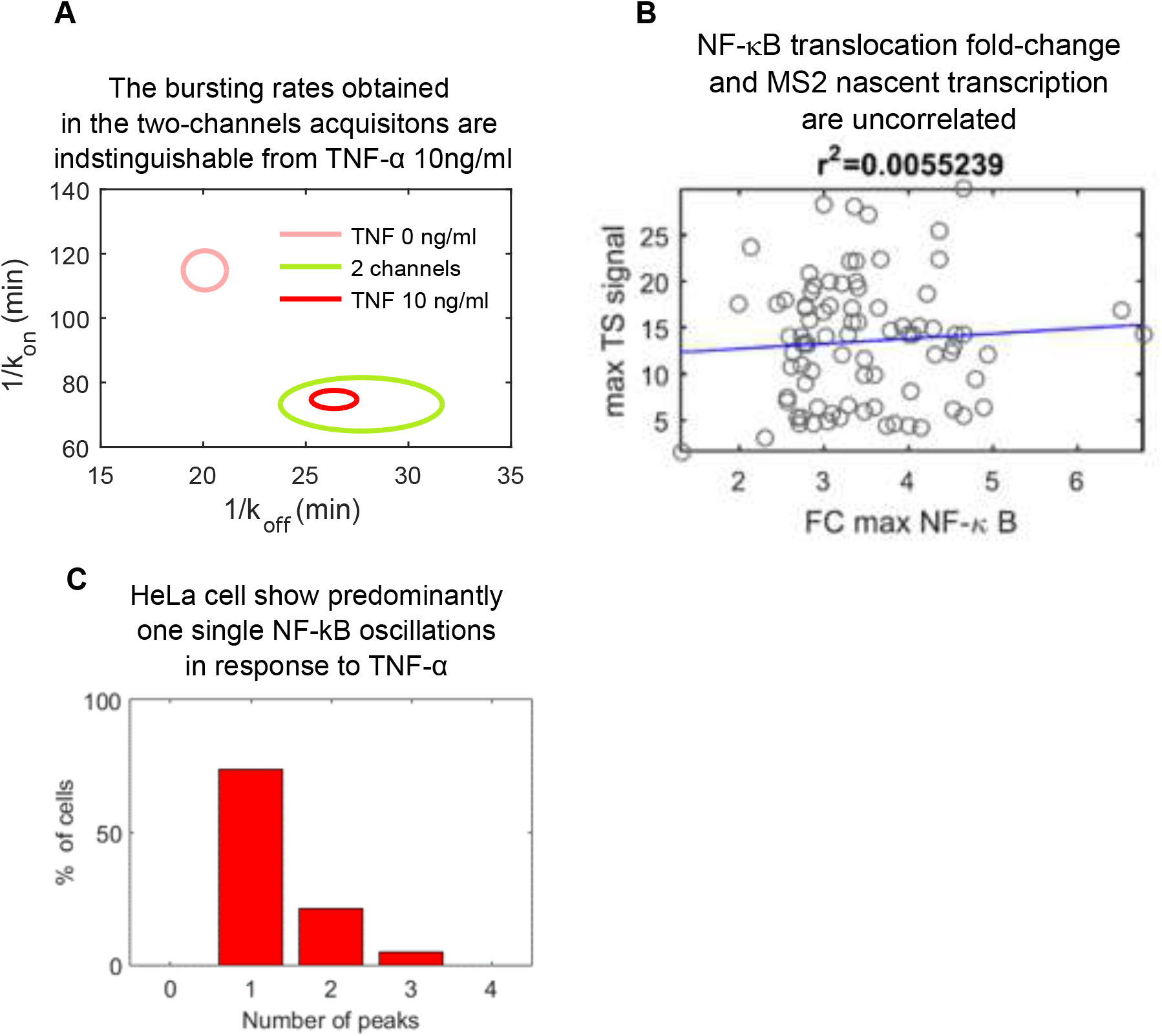
**A.** Parameters obtained by the stochastic fitting of the experiments obtained upon stimulation 10ng/ml TNF-α are identical for transfected and untransfected cells. **B**. Maximum TS signal against fold change nuclear NF-κB showing no correlation. **C.** Distribution of cells having different numbers of peaks of NF-κB activation. HeLa cells typically do not display oscillations, so cells with just one peak are the most frequents.

**Figure S6, related to Figure 6.**
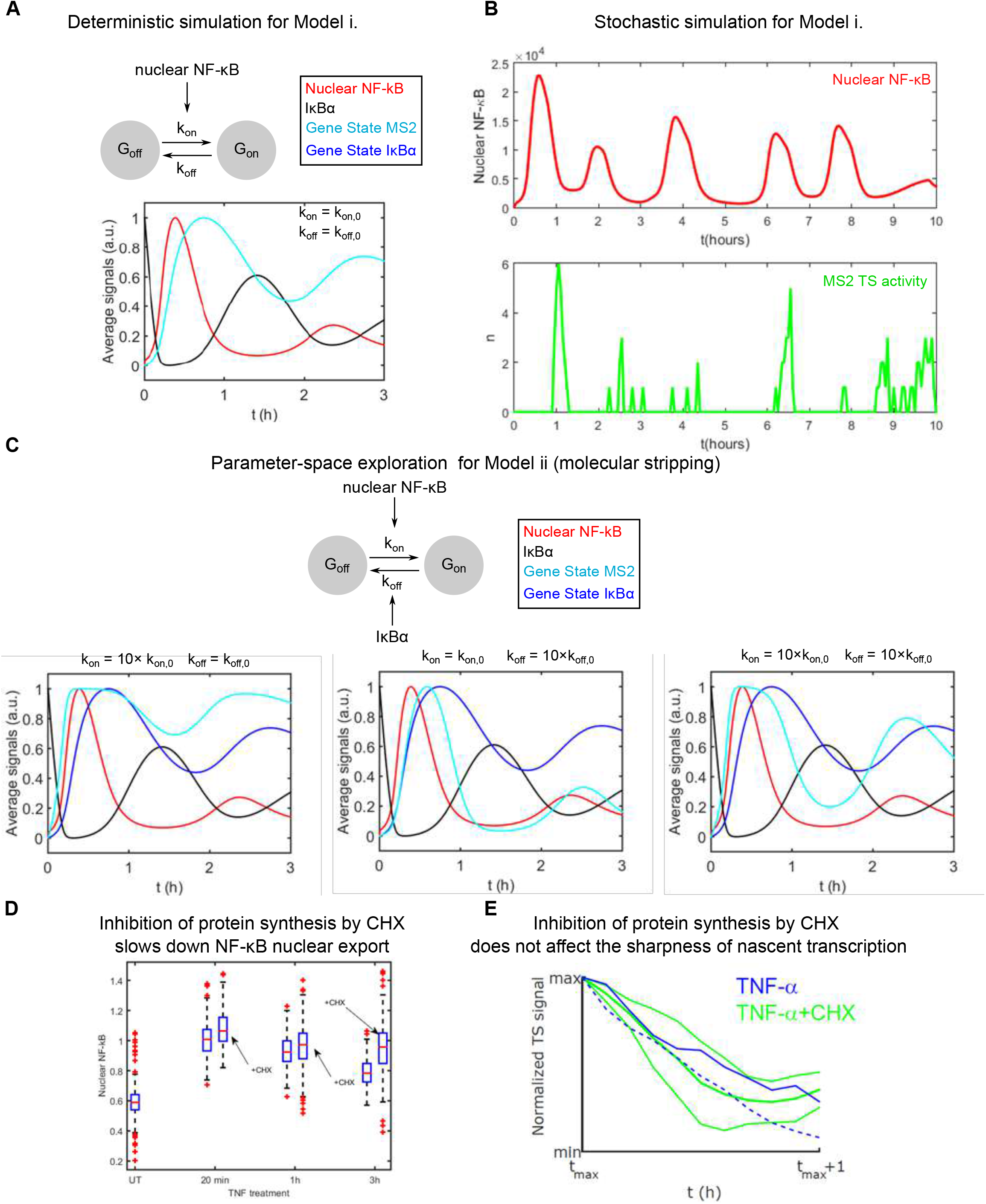
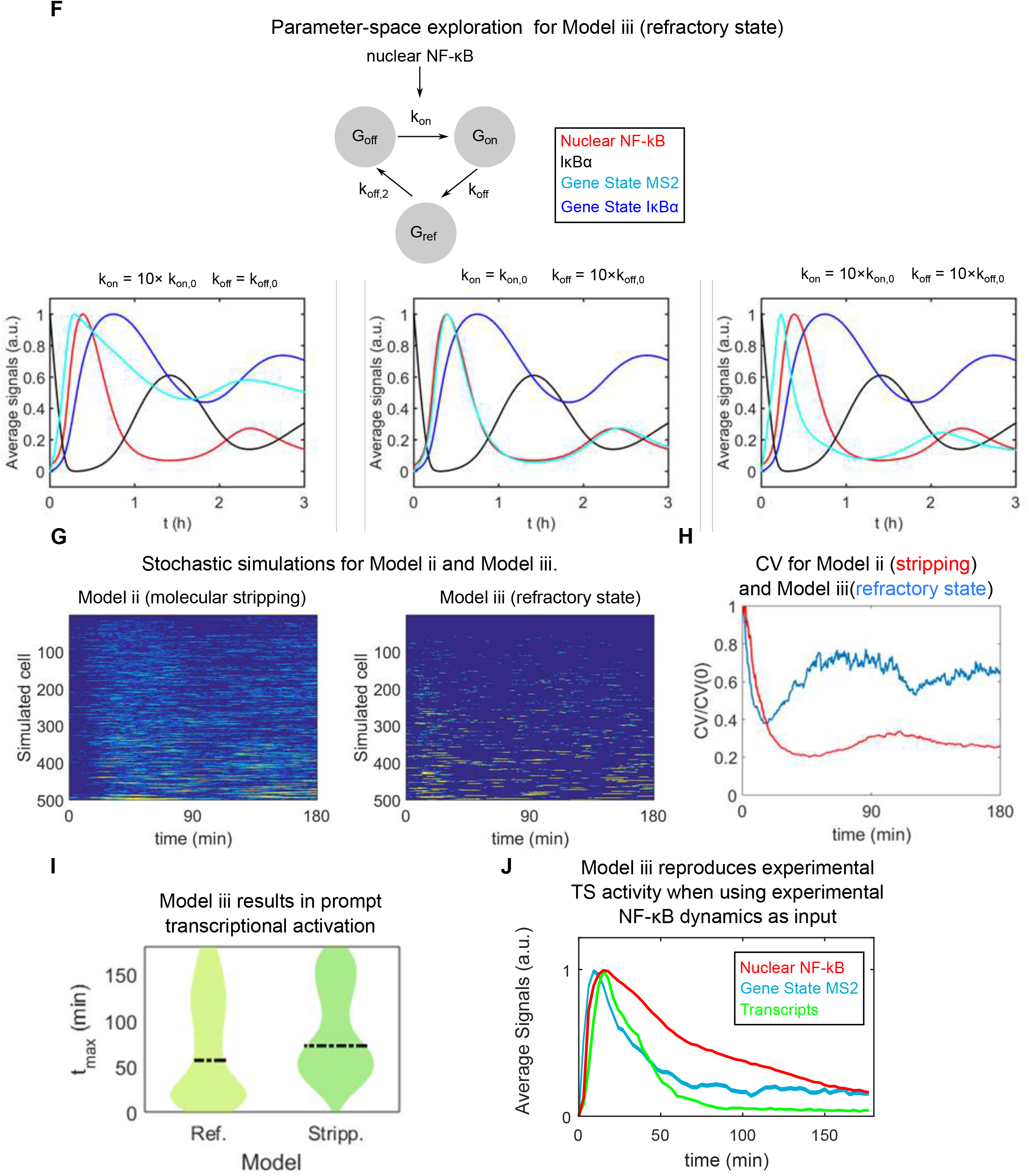
**A.** Deterministic simulations of NF-kB activation, inhibitor IκBα gene concentration, IκBα gene activity and target gene activity for the reference values used in the simulations. **B.** Stochastic simulations of the oscillations and bursts of nascent transcription obtained for the stochastic mathematical model, see Supplementary Methods. **C**. Simulations of the dynamics of the target gene activity obtained when considering the model with stripping, showing that prompt and sharp response are only obtained for parameter values out of the range considered. **D**. Quantification of the activation of NF-κB alone and in combination with cycloheximide (CHX): the latter gives rise to a more sustained activation. **E.** Decay of the maximum TS signal with CHX (blue lines, two independent experiments) and in absence of CHX (lines indicate mean and standard deviation of 4 experiments). CHX effect in the decay is negligible, further highlighting that stripping is not a plausible mechanism for the sharpness of the transcriptional response. **F.** Simulations of the dynamics of the target gene activity obtained when considering the model with a refractory state, showing that a sharp and prompt response can be obtained within the parameter range considered. **G.** Stochastic simulations performed with the model in which the gene is under stripping mechanism and by a refractory state, showing that the latter is able to reproduce the dynamics observed in the experiments. **H**. The coefficient of variation of simulated n(*t*) for the same two models, showing that he model including a gene refractory state and an NF-κB mediated activation mirrors the behavior observed in the experiments. **I.** The distribution of the timing of the maxima tmax for the simulations of the two models. The one for the gene refractory state (Ref) is reminiscent to the one obtained experimentally, including the fraction of first responders. **J.** Average simulated nascent transcriptional response and inferred gene state obtained using the NF-κB nuclear localization dynamics of single cells as driver of gene activation of our Model iii. with a refractory gene state. The prompt and sharp transcriptional response is reproduced.

## Notes

### Competing Interest Statement

The authors have declared no competing interest.

